# Glycerol-3-phosphate activates ChREBP, FGF21 transcription and lipogenesis in Citrin Deficiency

**DOI:** 10.1101/2024.12.27.630525

**Authors:** Vinod Tiwari, Byungchang Jin, Olivia Sun, Edwin D.J. Lopez Gonzalez, Min-Hsuan Chen, Xiwei Wu, Hardik Shah, Andrew Zhang, Mark A. Herman, Cassandra N. Spracklen, Russell P. Goodman, Charles Brenner

**Affiliations:** Beckman Research Institute of City of Hope; Duarte, USA; Liver Center and Endocrine Unit, Massachusetts General Hospital; Boston, USA; Comprehensive Cancer Center, University of Chicago; Chicago, USA; Baylor College of Medicine; Houston, USA; Department of Biostatistics and Epidemiology, University of Massachusetts; Amherst, USA

## Abstract

Citrin Deficiency (CD) is caused by inactivation of SLC25A13, a mitochondrial membrane protein required to move electrons from cytosolic NADH to the mitochondrial matrix in hepatocytes. People with CD do not like sweets. We discovered that SLC25A13 loss causes accumulation of glycerol-3-phosphate (G3P), which activates carbohydrate response element binding protein (ChREBP) to transcribe FGF21, which acts in the brain to restrain intake of sweets and alcohol, and to transcribe key genes of *de novo* lipogenesis. Mouse and human data establish G3P-ChREBP as a new mechanistic component of the Randle Cycle that contributes to metabolic dysfunction-associated steatotic liver disease (MASLD) and forms part of a system that communicates metabolic states from liver to brain in a manner that alters food and alcohol choices. The data provide a framework for understanding FGF21 induction in varied conditions, suggest ways to develop FGF21-inducing drugs, and drug candidates for both lean MASLD and support of urea cycle function in CD.

## Main

Citrin Deficiency (CD) is an autosomal recessive disease caused by mutation of the Citrin gene, *SLC25A13*, which encodes a mitochondrial membrane protein highly expressed in hepatocytes with a key role in moving high energy electrons from cytosol to the mitochondrial matrix^1,2^.

Infants with CD can be diagnosed in their first month by presenting with jaundice and elevated circulation of ammonia, citrulline and arginine, resembling a urea cycle disorder^3^, coincident with elevated lactate, resembling a mitochondrial disease^4^. Though CD is pan-ethnic^5^, it is most frequently diagnosed in the Far East. Data indicate a pathological allele frequency of up to 1 in 28 in Southern China, 1 in 45 across other parts of China^6^ and 1 in 50-100 elsewhere in the Far East^7^. CD is underdiagnosed outside of the Far East^8^, with a global disease burden that remains not well calculated and with unknown effects for *SLC25A13* mutation carriers.

By weight, carbohydrates constitute the largest class of macronutrient in human breast milk^9^ such that infant livers are primed to use carbohydrates as fuel, which requires nicotinamide adenine dinucleotide (NAD) coenzymes in the hydride-accepting NAD^+^ form in both the cytosol and the mitochondrial matrix^10^. As shown in Fig. 1A, glycolysis yields pyruvate and NADH in the cytosol. While pyruvate can be transported to the mitochondrial matrix for further oxidation^11^, cytosolic NADH does not cross the mitochondrial membrane^12^. Rather, the high energy electrons—termed reducing equivalents—picked up by NAD^+^ are moved to mitochondrial electron transport by two major NADH shuttle systems, the malate-aspartate (Asp) shuttle (MAS)^13^ and the glycerol-3-phosphate (G3P) dehydrogenase shuttle (GPDS)^14^. In hepatocytes, SLC25A13 is the component of the MAS that mediates Asp entry into cytosol in exchange for glutamate (Glu) transport into the mitochondrial matrix^15^. Another Glu/Asp antiporter encoded by SLC25A12 has higher expression in brain and muscle—its expression in hepatocytes is considered a primary mechanism of disease modification and/or escape in CD^16^.

**Fig. 1.**
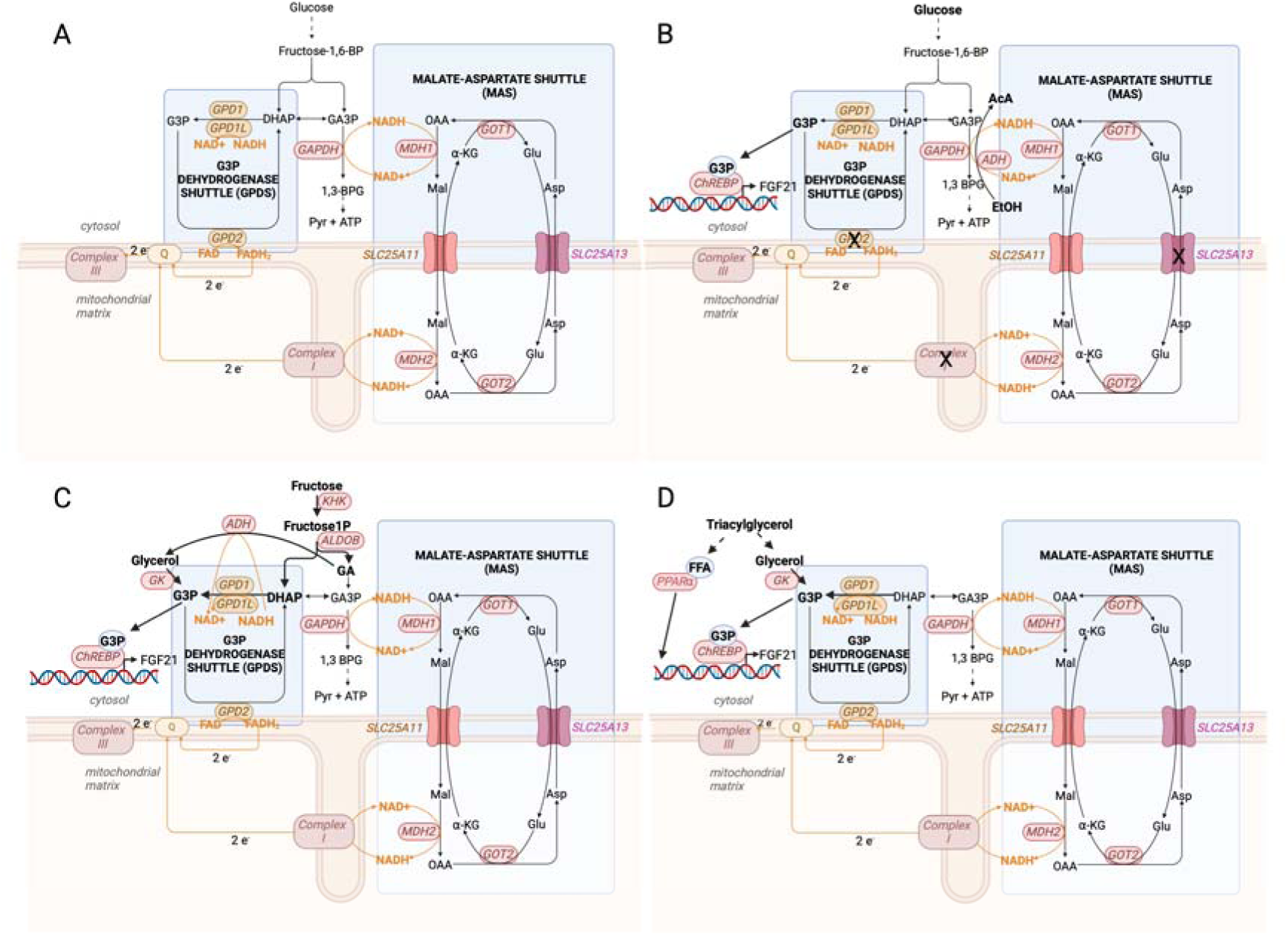
Inductive reasoning of a G3P-ChREBP-FGF21 transcription system in CD hepatocytes. FGF21 is induced in a wide variety of conditions that have eluded a unified theory of induction. The following diagrams of metabolic flow in hepatocytes as affected by conditions of metabolic stress led us to propose G3P as the activator of ChREBP that can resolve paradoxes of FGF21. (A) Metabolic flow in unperturbed hepatocytes is facilitated by two NADH shuttles without induction of FGF21. The MAS^1^ facilitates transfer of reducing equivalents from cytosolic NADH to oxaloacetate, transiently forming malate, which is reoxidized at complex I in the mitochondrial matrix. The GPDS^13^ facilitates transfer of reducing equivalents from cytosolic NADH to dihydroxyacetone phosphate (DHAP), transiently forming glycerol-3-phosphate (G3P) with reducing equivalents transferred to FAD, forming FADH_2_, which is reoxidized with electron transfer to coenzyme Q in the mitochondrial electron transfer chain (METC). (B) With either mitochondrial insufficiency, disruption of the MAS, disruption of the GPDS, ethanol metabolism or elevated glucose, cytosolic NADH is expected to rise, which would be expected to cause accumulation of G3P, the proposed activator of ChREBP, driving FGF21 expression. (C) Fructolysis is predicted to elevate G3P with one G3P equivalent formed from DHAP and another formed from glyceraldehyde (GA), via function of alcohol dehydrogenase (ADH) and glycerol kinase^97^. The resulting G3P is proposed as the activating ligand for ChREBP activation and FGF21 expression. (D) Triglyceride lipolysis is expected to produce glycerol and free fatty acids. Glycerol release and conversion to G3P is proposed as the activating ligand of ChREBP, which would co-operate with free fatty acid-activated PPARα to drive transcription of FGF21.

The resemblance of CD to a mitochondrial disease can be explained by failure of CD hepatocytes to obtain mitochondrial energy from complete oxidation of carbohydrates, with elevated lactate being an expected outcome of elevated cytosolic NADH. The resemblance of CD to a urea cycle disorder can be explained by a deficiency in hepatocytosolic Asp, which is required for the citrulline-consuming step of the urea cycle^5^. After diagnosis, children with CD, who are classified as cases of neonatal intrahepatic cholestasis caused by CD (NICCD)^2,3^ are managed nutritionally with a diet in which carbohydrates are largely replaced by medium chain triglycerides (MCTs)^5^. In most cases, NICCD goes into remission and people with CD are able to grow and lead relatively normal lives, though they do not like sweets^3,17^ and are prone to metabolic dysfunction-associated steatotic liver disease (MASLD) despite a lean body mass^18^. In other cases, there is a failure to thrive with dyslipidemia caused by CD (FTTDCD). In adulthood, CD symptomology can reactivate with diagnoses of adult-onset type II citrullinemia (CTLN2) or adult-onset CD (AACD), which are characterized by hyperammonemia, MASLD, pancreatitis and neuropsychiatric complications^2,3^. Conventional nutritional management of hyperammonemia, *i.e.*, a lower protein, higher carbohydrate diet, does not benefit patients with CD^19^.

When *Slc25a13* was knocked out in mice, there was no clear physiological phenotype^20^. Reasoning that the GPDS is more highly expressed in mouse liver than in human, researchers proceeded to inactivate GPDS component *Gpd2*. Mice homozygous for disruption of *Slc25a13* and *Gpd2* formed an excellent CD model, exhibiting hyperlactatemia, hyperammonemia, hyperargininemia, hypercitrullinemia and MASLD^14^. When provided a choice between water and saccharine, *Slc25a13* -/- *Gpd2* -/- mice behaved as wild-type, choosing saccharine by a wide margin, indicating that there is no defect in detection of or desire for sweet taste in naïve mice with inactivation of the two NADH shuttle systems^21^. However, when provided with a choice between sucrose and water, *Slc25a13* -/- *Gpd2* -/- double mutants fail to prefer sucrose, suggesting that incomplete carbohydrate oxidation is required to produce the carbohydrate- aversive phenotype. Though wild-type mice consumed greater than 15 grams of sucrose or 3 grams of ethanol or 4 grams of glycerol per 25 gram mouse per day, *Slc25a13* -/- *Gpd2* -/- double mutants had impaired preferences for sucrose, ethanol and glycerol when given a choice between water and these energy-containing liquids^21^. Single homozygous mutants in the MAS component *Slc25a13* and the GPDS component *Gpd2* generally had intermediate phenotypes. Metabolomic analysis showed accumulation of glycerol and G3P in *Slc25a13* -/- *Gpd2* -/- double mutants^21^. These data suggested that loss of the MAS and GPDS coupled with provision of specific macronutrients result in a type of metabolic stress that leads to a behavioral change to avoid these compounds.

Fibroblast growth factor 21 (FGF21) is a secreted polypeptide made in liver, adipose and muscle in response to a wide variety of stress conditions^22^. FGF21 has multiple sites of action including specific β-klotho-expressing regions of the brain, where FGF21 functions to restrain intake of sweets^23^ and ethanol^24^, and the periphery, where it increases energy expenditure^25^ and body temperature^26^. FGF21 was first termed a starvation hormone because it is released into circulation by the liver in response to fasting^27,28^ and ketogenic diet (KD)^29^ in mice. Though these conditions result in release of free fatty acids, which activate the PPARα transcriptional coactivator-binding site^30^ in the *FGF21* promoter, deletion of PPARα from mouse liver did not completely eliminate induction of FGF21 by KD^29^. The literature on FGF21 induction is considered paradoxical^31^ because FGF21 is not only induced by fasting and the near absence of dietary carbohydrates but also by provision of simple carbohydrates^32–34^, particularly fructose^35,36^. Moreover, in addition to being induced by fasting^27,28,37^ and exercise^38,39^, FGF21 is induced by refeeding^40^, obesity^41^, type 2 diabetes^42^ and mitochondrial disease^43^. It has also been shown that FGF21 is induced by a loss-of-function variant of glucokinase regulator *GCKR*^44^ and ethanol^45^ via reductive stress, i.e., conditions that increases the NADH/NAD^+^ ratio in hepatocytes^46^, and by protein restriction^47^.

We considered whether mitochondrial disease, NADH shuttle disruption, ethanol, fructolysis and the conditions involving lipolysis, i.e., fasting and exercise, might produce a common metabolite that would activate FGF21 transcription. As shown in Fig. 1B, we reasoned that mitochondrial disease, NADH shuttle disruption, elevated glycemia and/or ethanol metabolism would elevate cytosolic NADH, leading to a buildup of G3P at the expense of dihydroxyacetone phosphate (DHAP). As shown in Fig. 1C, we reasoned that fructolysis could also lead to a buildup of G3P with G3P formed from DHAP and glyceraldehyde. Finally, as shown in Fig. 1D, we reasoned that triglyceride lipolysis would produce glycerol and, consequently, G3P due to the activity of glycerol kinase.

Two of the key transcription factors acting within the *FGF21* promoter are PPARα and carbohydrate response element binding protein (ChREBP)^31,34^. While it is clear that PPARα strongly contributes to turning on FGF21 in conditions of fasting^27–29^ and that FGF21 is both upstream and downstream of PPARα in the adaptive response to starvation^28^, the PPARα- independent component of FGF21 induction in mice fed a KD^29^ suggested the action of a different metabolite-sensing factor.

ChREBP, encoded by the *MLXIPL* gene, is a member of the MYC and MAX superfamily of heterodimerizing helix-loop-helix transcription factors that is abundantly expressed in liver^48–50^. As a heterodimer with MAX-like protein X (MLX) and at elevated levels of glucose (Glc) metabolites, ChREBP activates transcription of target promoters containing carbohydrate response elements (ChoREs)^51^. Well characterized ChoREs drive ChREBP-dependent transcription of ChREBPβ—a shorter, carbohydrate-induced form of ChREBP^52^, liver pyruvate kinase (*PKLR*)^53^, *FGF21*^31^ and other genes that are important for carbohydrate adaptations including those for *de novo* lipogenesis^51^ such as Ac-coA lysase (ACLY), Ac-coA carboxylase (*ACACA*), and fatty acid synthetase (*FASN*).

In the extensive literature on ChREBP^48–50^, researchers have reported that ChREBP is activated by a Glc metabolite that engages the N-terminal Glc-sensing module (GSM)^54^, which is conserved between ChREBP and the MondoA transcription factor^55^. The specific identity of this metabolite remains less clear, however, as evidence has been presented for glucose-6-phosphate (G6P)^55,56^, fructose-2-6-bisphosphate (F2,6BP)^57^ and xylulose-5-phosphate X5P^58^. It is difficult to distinguish between these proposed mechanisms because typical ChREBP activation conditions involve a shift from low Glc to high Glc that would simultaneously elevate all proposed ChREBP GSM-activating ligands, and none of these metabolites have been shown directly to bind the GSM. Moreover, it is challenging to reconcile the previously proposed ChREBP ligands with data showing that glycerol treatment strongly activates hepatic ChREBP in vivo^59^.

Recent work showed that the key transcription factor for ethanol induction of FGF21 is ChREBP and that the ChREBP transcription program is downstream of an increase in the NADH/NAD^+^ ratio that occurred with a rise in a select group of metabolites that include G3P but not G6P or X5P^60^. We hypothesize that CD patients and mouse models have a sweet and ethanol aversive phenotype because their livers activate a G3P-ChREBP-FGF21 transcription program (Fig. 1B). Moreover, we suggest that this mechanism resolves what have been considered paradoxical aspects of FGF21 induction because fasting and refeeding, ketogenic diet and simple carbohydrates, mitochondrial disease, ethanol and fructose all have direct paths to producing G3P (Fig. 1B-D).

Here we show that G3P accumulates in the liver of the mouse model of CD, and that ChREBP is activated in this model with transcription of FGF21 and activation of a lipogenic transcriptional program. Our data further show that G3P is a specific ligand of the ChREBP GSM, and suggest that features of the G3P-ChREBP activation mechanism can account for why fructose is more lipogenic than Glc, provide a unifying mechanism for nonalcoholic and alcoholic hepatic steatogenesis, resolve paradoxes of FGF21 expression, and explain key aspects of CD pathogenesis including lean MASLD, the favorable effects of MCTs, and severe urea cycle dysfunction. This work suggests drug targets for treatment of CD, lean MASLD and common MASLD, and also implicates CD mutation carriers, who number in the millions, as people with a distinct metabolic profile consistent with elevated circulation of FGF21.

## Results and Discussion

### Deletion of NADH shuttle systems and provision of glycerol cause increased FGF21 circulation in mice

Sperm from C57BL/6J mice of genotype *Slc25a13* -/- *Gpd2* -/-^21^ were used for *in vitro* fertilization of C57BL/6J females. Subsequent crosses generated male and female *Slc25a13 -/-* mice, *Gpd2 -/-* mice and, at much lower than Mendelian ratios, *Slc25a13* -/- *Gpd2* -/- mice.

Consistent with our first prediction, as shown in Fig. 2A, mice with deletion of either *Slc25a13* or *Gpd2* tended to have elevated circulating FGF21 while mice inactiveated for both NADH shuttle systems have ∼3-fold elevated FGF21. This result was significant for mice of both sexes (Fig. 2A), for male mice (Fig. 2B) and for females (Fig. 2C). Mice of each genotype with 5% (w/v) glycerol in their drinking water for 2 days had significantly elevated circulating FGF21 by virtue of glycerol exposure and the effects of genotype and glycerol were additive. The effect of glycerol on induction of FGF21 was highly significant in each genotype irrespective of sex (Fig. 2A) and in males (Fig. 2B). Further, *Slc25a13* -/- *Gpd2 -/-* females have significantly higher circulating FGF21 than wild-types or either single mutant upon exposure to glycerol (Fig. 2C). Thus, consistent with the hypothesis that sweet aversion in CD is due to a G3P-ChREBP-FGF21 induction program, the mouse model of CD overproduces FGF21, and superinduces FGF21 when exposed to glycerol.

**Fig. 2.**
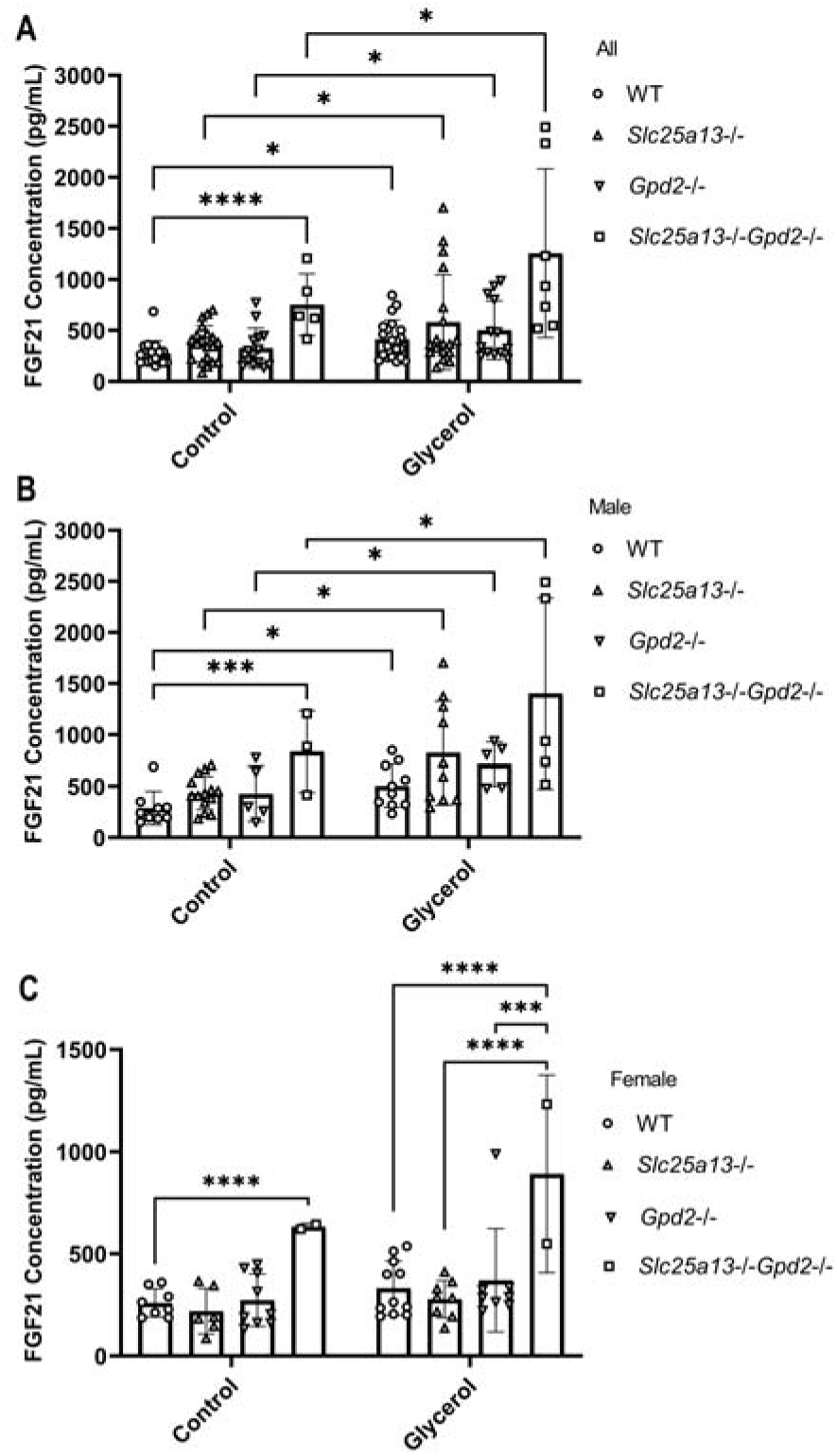
Inactivation of NADH shuttle systems and glycerol induce FGF21 circulation. (A) Serum FGF21 from mice of indicated genotypes were measured after two days with *ad libitim* access to food and water (wild type [o] n=17, *Slc25a13 -/-* [Δ] n=20, *Gpd2 -/-* [∇] n=14, *Slc25a - /- Gpd2 -/-* (□) n=5, or food and 5% (w/v) glycerol (wild type [o] n=21, *Slc25a13 -/-* [Δ] n=18, *Gpd2 -/-* [∇] n=13, *Slc25a -/- Gpd2 -/-* [□] n=7. The data show that across both sexes, *Slc25a -/- Gpd2 -/-* mice have elevated FGF21 and that each genotype has its FGF21 circulation elevated by glycerol. (B) Subgroup analysis of male mice of indicated genotypes with *ad libitim* access to food and water (wild type [o] n=9, *Slc25a13 -/-* [Δ] n=14, *Gpd2 -/-* [∇] n=5, *Slc25a -/- Gpd2 -/-* [□] n=3, or food and 5% (w/v) glycerol (wild type [o] n=10, *Slc25a13 -/-* [Δ] n=10, *Gpd2 -/-* [∇] n=5, *Slc25a -/- Gpd2 -/-* [□] n=5. The data show that male *Slc25a -/- Gpd2 -/-* mice have elevated FGF21 and that each genotype has its FGF21 circulation elevated by glycerol. (C) Subgroup analysis of female mice of indicated genotypes with *ad libitim* access to food and water (wild type [o] n=8, *Slc25a13* -/- [Δ] n=6, *Gpd2* -/- (∇) n=9, *Slc25a13* -/- *Gpd2* -/- (□) n=2}, or food and 5% (w/v) glycerol (wild type [o] n=11, *Slc25a13* -/- [Δ] n=8, *Gpd2* -/- [∇] n=8, *Slc25a13* -/- *Gpd2* -/- [□] n=2}. The data show that female *Slc25a -/- Gpd2 -/-* mice have elevated FGF21 with respect to wild-type at baseline. In addition, the data show that female *Slc25a -/- Gpd2 -/-* mice have elevated FGF21 on glycerol with respect to each of the other genotypes on glycerol. Significant differences between were calculated by applying two-way ANOVA and Tukey’s multiple comparisons test in which *p<0.05, **p<0.005, *** p<0.0005 and ****p<0.0001.

### Deletion of NADH shuttle systems and provision of glycerol activate hepatic ChREBP plus FGF21 and lipogenic transcription

To determine whether deletion of NADH shuttle systems and/or provision of glycerol resulted in a ChREBP transcriptional program, we harvested livers from 40 mice representing water and glycerol-exposed males of the four genotypes, prepared cDNA and performed bulk paired-end 150 base-pair RNA sequencing (RNAseq)^61^ using an Illumina NovaSeq X Plus sequencer at >20 million paired reads per sample. As shown in Fig. 3, the ChREBP transcriptional program is evident as judged by induction of well characterized ChREBP target genes including *ChREBPβ*, *Fgf21*, *Pklr* (encoding liver pyruvate kinase), *Khk, Aldob, Tkfc* (the three key enzymes of fructolysis), *Gpi1, Pgd* (phosphoglucose isomerase and 6-phosphogluconate dehydrogenase), *Fasn, Elovl6, Agpat2* (key enzymes for triglyceride synthesis), and *Tm6sf2* (very low density lipoprotein synthesis factor).

**Fig. 3.**
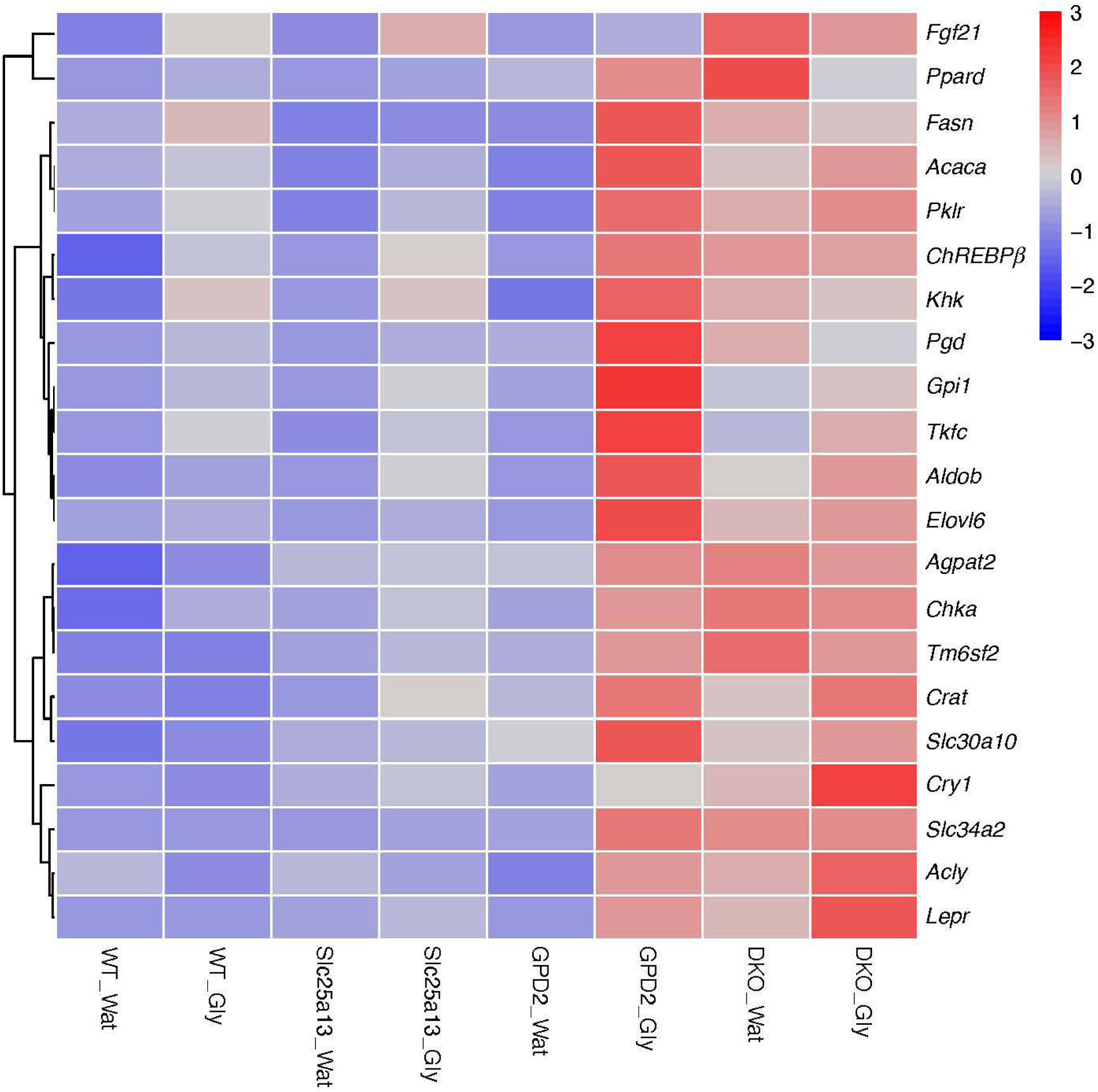
Inactivation of NADH shuttle systems and glycerol drive a ChREBP transcription program. Hierarchical clustering of mean liver gene expression levels (fragments per kilobase of transcript per million mapped reads, FPKM) of select ChREBP target genes across four genotypes of mice exposed to either water (Wat) or glycerol (Gly).

Many of the genes of *de novo* lipogenesis are dually activated by ChREBP and sterol response element binding protein 1c (SREBP-1c)^62^, encoded by the *Srebf1* gene. To test whether the mouse model of CD dysregulates either of these transcription factors and to examine the relationship with *Fgf21* expression, we calculated the levels of specific transcripts using RSEM^63^. As shown in Table 1, *Srebf1* expression was unaffected by deletion of *Slc25a13* and/or *Gpd2* and was unaffected by addition of glycerol. the *ChREBPβ* transcript was induced by about 2.4-fold by glycerol in the wild-type strain, about 4.5-fold by loss of either of the NADH shuttle genes, and nearly 13-fold by inactivation of both genes. The *ChREBPα* transcript was largely unaffected by genotype but was induced ∼2-3 fold by glycerol in wild type and single mutant strains. In the double mutant, which has the highest level of circulating FGF21 and the highest level of *ChREBPβ* transcript, there was no further induction of of *ChREBP* transcripts by glycerol. Consistent with the effect of FGF21 on feeding behaviors, at the mRNA level, glycerol induced *Fgf21* to a greater degree in the wild-type and *Slc25a13 -/-* strains in which basal *Fgf21* expression was lowest and induced *Fgf21* the least in the *Gpd2 -/-* and *Slc25a13 -/- Gpd2 -/-* strains in which basal *Fgf21* expression was highest. As the experiment was performed with 2 days of *<u>ad libitim</u>* access to 5% glycerol as the water source and it has been shown that the CD mouse model has a sucrose, ethanol and glycerol-aversive phenotype^21^, one would expect that the high basal levels of FGF21 in the *Slc25a13 -/- Gpd2 -/-* strain significantly limit their glycerol intake and further increases in *Fgf21* mRNA expression.

**Table 1.**
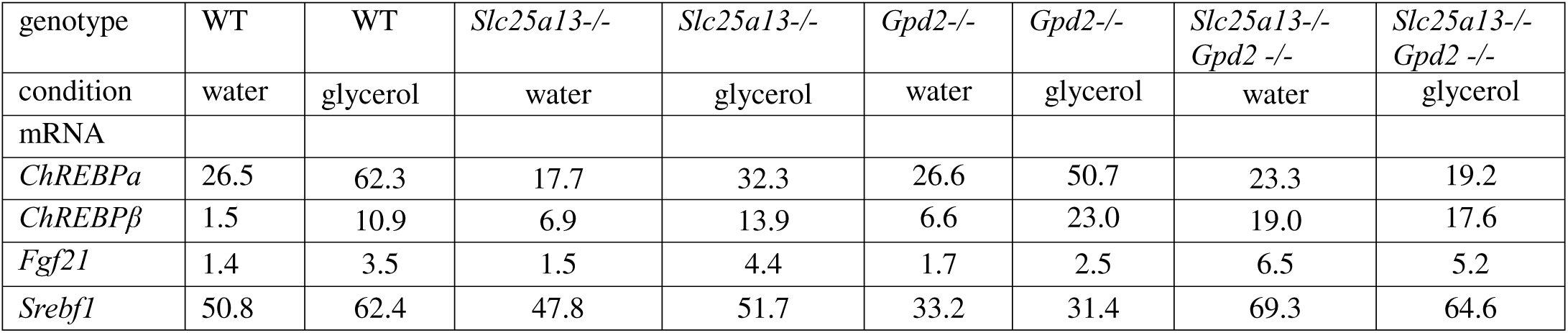
Hepatic gene expression (mean FPKM) with NADH shuttle gene deletion and/or glycerol.

### Deletion of NADH shuttle systems result in accumulation of hepatic G3P

We performed targeted metabolomic analysis of liver extracts from male mice of the four genotypes to determine whether G3P or compounds previously termed ChREBP activators accumulated as a function of NADH shuttle disruption. As shown in Fig. 4A, G3P can be quantified at ∼ 1 mM in liver from wild-type mice. Deletion of *Slc25a13*, *Gpd2* or both NADH shuttle genes gave levels of hepatic G3P that tended to be higher than levels of G3P in wild-type mouse livers. Similarly, the glycerol-exposed mouse livers of each of the four genotypes had levels of G3P that tended to be higher. On the basis that the ChREBPβ transcript is induced in *Slc25a13 -/-*, *Gpd2 -/-* and *Slc25a13 -/- Gpd2 -/-* mouse livers, we compared G3P levels in wild-type versus all mutant livers and observed a significant increase in G3P from 1.09 +/- 0.26 mM to 1.98 +/- 0.59 mM. As glucose-6-phosphate (G6P)^55,56^, fructose-2-6-bisphosphate (F2,6BP)^57^ and xylulose-5-phosphate X5P^58^ have been previously proposed to be activating ligands of ChREBP, we tested whether hexose phosphates or pentose phosphates were altered by conditions that activate ChREBP, namely glycerol and deletion of NADH shuttle genes. As shown in Fig. 4C and 4D, hepatic concentrations of hexose phosphates were not altered by genotype and/or addition of glycerol. As shown in Fig. 4E and 4F, relative levels of pentose phosphates were also unaffected. Thus, consistent with the proposed G3P-ChREBP-FGF21 activation mechanism in Fig. 1B and the observation that *Slc25a13 -/- Gpd2 -/-* mice experiencing sweet and ethanol aversive behaviors have elevated hepatic G3P^21^, we show that deletion of NADH shuttle systems is sufficient specifically to elevate the proposed ChREBP-activating ligand.

**Fig. 4.**
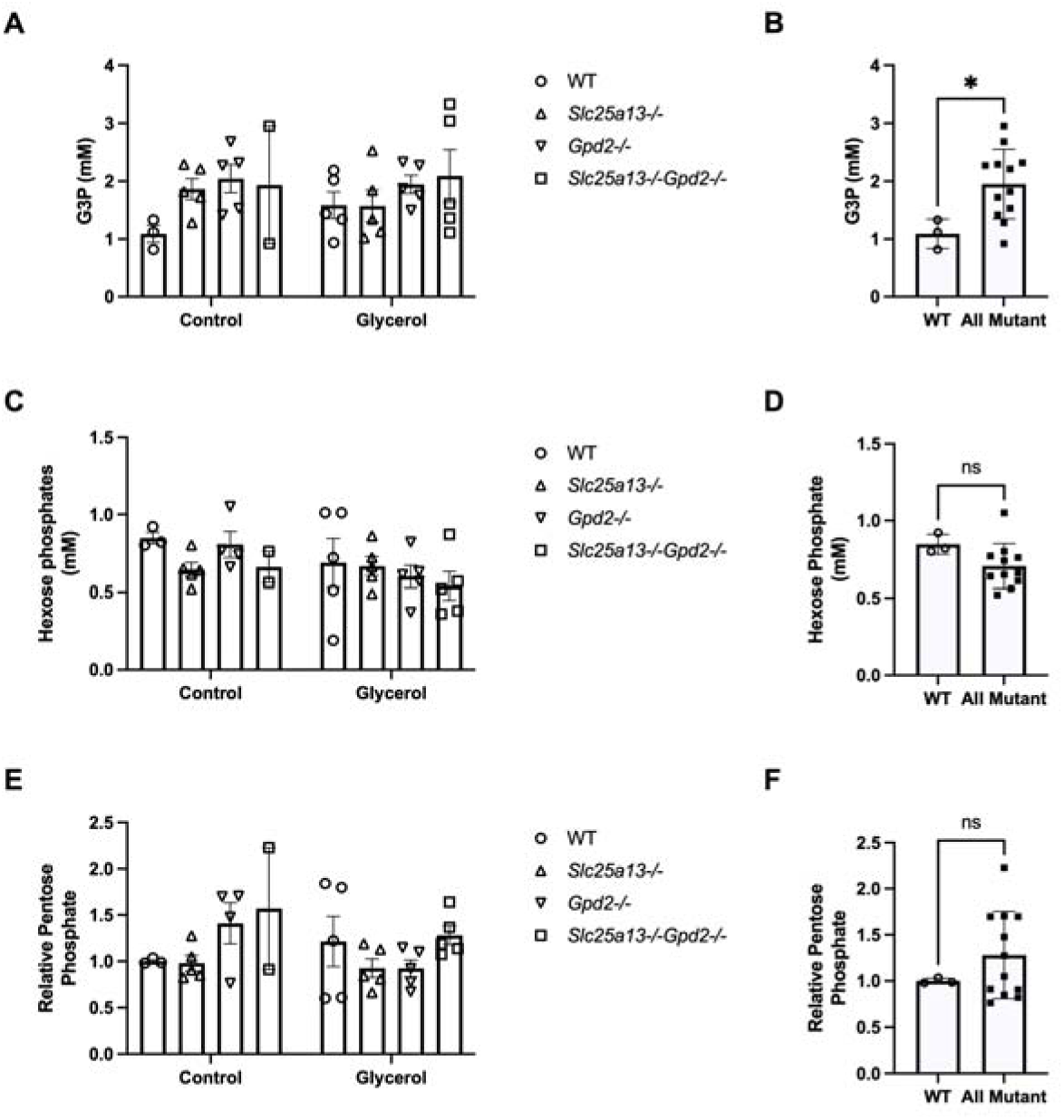
NADH shuttle disruption elevates hepatic G3P. We performed quantitative metabolomics for hepatic G3P (A and B) and hexose phosphates (C and D), and qualitative metabolomics for hepatic levels of pentose phosphates (E and F). Numbers of mice analyzed were wild-type (o, n=3), *Slc25a13 -/-* (Δ, n=5), *Gpd2* (∇, n=5) and *Slc25a13 -/- Gpd2 -/-* (□; n=2) in mice with two days of *ad libitim* access to food and water. Numbers of mice analyzed were wild-type (o, n=5), *Slc25a13 -/-* (Δ, n=5), *Gpd2* (∇, n=5) and *Slc25a13 -/- Gpd2 -/-* (□; n=5) with two days of *ad libitim* access to food and 5% w/v glycerol. In (A), (C) and (F), all mouse genotypes were analyzed separately. In, (B), (D) and (F), the three mutant genotypes were pooled to compare wild-type (o, n=3) to the three genotypes containing one or both NADH shuttle systems disrupted (▄, n=12) with exposure to water. In (A), the data show that each mutant genotype tends to have higher G3P than wild-type and that glycerol tends to elevate G3P with respect to water. In (B), the data show that deletion of one or both NADH shuttle systems significantly elevates G3P (p=0.03 by unpaired t-test, t = 2.368, degrees of freedom =13. (C and D) No differences were observed in hexose phosphate levels across the four genotypes and two conditions. (E and F) No differences were observed in hexose phosphate levels across the four genotypes and two conditions. *p<0.05.

### Genetic manipulation of G3P drives ChREBP activation in a reconstituted system

HEK293T cells do not express ChREBP and show poor expression of MLX, thereby allowing reconstitution of condition-dependent, ChREBP-MLX-dependent, ChoRE-dependent transcription^60^. Robust ChoRE-luciferase activity in HEK293T cells depends on introduction of both ChREBPα and MLX. Reconstituted transcriptional activity is depressed by expression of *Lb*NOX, which lowers the NADH/NAD^+^ ratio, and increased by expression of *E. coli* soluble transhydrogenase (*Ec*STH), an enzyme that uses reducing equivalents from NADPH to elevate the cytosolic NADH/NAD^+^ ratio^60^. In prior work, this system was used to show that three metabolites, namely G3P, glyceraldehyde-3-phosphate (GA3P) and fructose-1,6-bisphosphate (F1,6BP), were highly correlated with ChREBPα activation, but that levels of G6P and X5P were uncorrelated with ChREBP transcriptional activity^60^. To distinguish between potential ChREBPα-activating ligands, we introduced glycerol kinase (*GK*), *GPD1* and *GAPDH* genes into the reconstituted ChREBPα-MLX HEK293T system alongside green fluorescent protein (GFP), *Lb*NOX and *Ec*STH as inactive, ChoRE luciferase-depressing and ChoRE luciferase- activating controls, respectively. Though *GAPDH* had a minor ChoRE-luciferase activity- depressing effect and *GK* was without effect potentially due to the absence of a supply of glycerol, *GPD1* strongly increased ChoRE-luciferase activity (Fig. 5A). Relative quantification of 137 metabolites showed that in these cells, *GPD1* strongly depressed levels of GA3P and DHAP, while elevating G3P. The correlation coefficient (CC) for luciferase activity with relative G3P in the six resulting HEK293T transfected cell lines was 0.96 (Fig. 5B), exceeding all other compounds tested (Supplementary Data). G6P (Fig. 5C), which has been considered a candidate ChREBP-activating ligand^55,56^, was uncorrelated with ChoRE-luciferase activity (CC = -0.11). GA3P (Fig. 5D), which had appeared to be correlated to ChORE-luciferase activity in prior work^60^ became uncorrelated with inclusion of the effects of *GK*, *GPD1* and *GAPDH* (CC = 0.18).

**Fig. 5.**
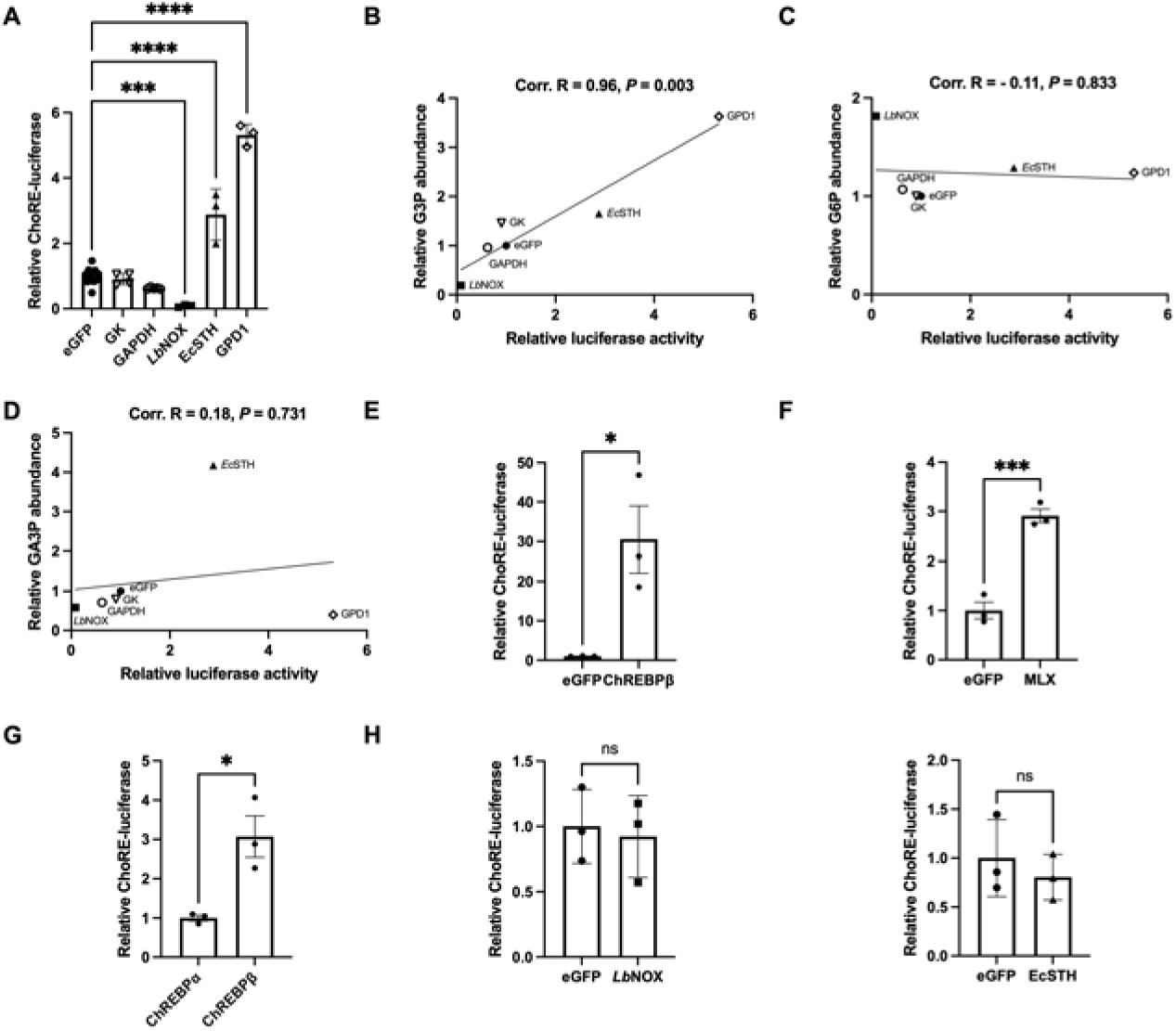
The ChREBPα-specific isoform of ChREBP is activated in a manner that is coincident with G3P accumulation. (A) HEK293T cells co-transfected with ChREBPα, MLX, ChORE-luciferase and the indicated genetic constructs were assessed for ChORE-luciferase activity. The data show that *LbNOX* depresses, *EcSTH* increases and *GPD1* greatly increases ChORE-luciferase activity. Significant differences between were calculated by applying one-way ANOVA and Dunnett’s multiple comparisons test in which *** p<0.0005 and ****p<0.0001. (B- D) The relative levels of 137 metabolites were determined by LC-MS. By plotting ChoRE- luciferase activity against each metabolite we show that G3P is highly correlated (coorelation coefficient = 0.96) with ChREBPα-dependent ChoRE-luciferase activity (B) while (C) G6P and (C) are not. The P values tested the hypothesis of a nonzero slope. (E) HEK293T cells co- transfected with ChREBPβ or eGFP plus ChoRE-luciferase were assessed for ChORE-luciferase activity. (F) HEK293T cells co-transfected with ChREBPβ with MLX or eGFP plus ChORE- luciferase were assessed for ChORE-luciferase activity. (G) HEK293T cells co-transfected with MLX plus ChORE-luciferase plus either ChREBPα or ChREBPβ were assessed for ChORE- luciferase activity. Finally, HEK293T cells co-transfected with ChREBPβ, MLX and ChORE- luciferase plus were assayed for ChoRE-luciferase activity with either (H) eGFP or *Lb*NOX or (I) eGFP or *Ec*STH. Data in (E-I) show that ChREBPβ has significant ChoRE-luciferase activity in HEK293T cells (E) that is further boosted by MLX transfection (F) and that the ChoRE- luciferase activity of ChREBPβ-MLX exceeds that of ChREBPα-MLX (G). However, in contrast to the ability of *Lb*NOX to depress and *Ec*STH to increase ChoRE-luciferase activity of ChREBPα-MLX (A), ChREBPβ-MLX was not regulated by either *Lb*NOX (H) or *Ec*STH (I), thereby mapping modulation of ChREBPα to the N-terminal GSM domain. Significant differences were calculated with unpaired t-tests in which *p<0.05 and *** p<0.0005.

To further map the site of G3P activity, we used the HEK293T system to characterize the sensitivity of ChREBPβ-MLX to altered levels of metabolites. As shown in Fig. 5E-G, ChREBPβ—the form of ChREBP without the N-terminal GSM^52^—was introduced into HEK293T cells and shown to induce ChoRE-luciferase in a manner that depends on MLX cotransfection. Moreover, ChoRE-luciferase activation from ChREBPβ-MLX was ∼3-fold more potent than that from ChREBPα-MLX (Fig. 5H). However, when ChREBPβ-MLX-dependent ChORE-luciferase was challenged by *Lb*NOX (Fig. 5I) and *Ec*STH (Fig. 5J) expression, there was no modulation of ChREBP transcriptional activity^60^. These data indicate that the metabolite that responds to an elevated NADH/NAD^+^ ratio thereby driving ChREBP-MLX transcription^60^ acts on the ChREBPα-specific N-terminus and is fully correlated with accumulation of G3P.

### The GSM domain of ChREBP is a G3P-sensing module

Various constructs have been used to obtain structural or biophysical data on the GSM of ChREBP. Notably, a construct from residue 1 to 250 of mouse ChREBP was purified as a His-tagged protein in *E. coli* for structural characterization. However, this molecule was found to have been proteolyzed to a fragment of the GSM from residue 81 to 196 and, when mixed with 14-3-3β protein for structural characterization, the full length of 14-3-3β and only residues 117 to 137 of ChREBP were structured^64^. Based on knowledge that both ChREBP and the homologous MondoA are responsive to Glc metabolites^54,65^, we performed a careful multiple sequence alignment of ChREBP and MondoA and chose to define residues 43 to 307 of mouse ChREBP (ChREBP43-307) as a candidate stable, globular GSM.

After expression of His-tagged ChREBP43-307 in *E. coli* and purification by nickel nitrilotriacetic acid affinity chromatography, we characterized ligand binding by isothermal titration calorimetry^66^. As shown in Table 2, we tested Glc, G6P, F6P, F1,6BP, GA3P, DHAP, G3P and X5P for binding and were able to detect saturable binding with each ligand. However, the *K*_d_ values for all but two ligands were greater than 130 μM. G6P, which has been considered a candidate GSM ligand^55,56^ but is uncorrelated with ChREBP transcriptional activation^60^ (Fig. 5C), showed half-saturated binding at 64.3 +/- 13.5 μM, suggesting an association that could be displaced by a higher affinity ligand whose abundance is sensitive to conditions that activate ChREBP. Indeed, G3P, which is greatly increased by deletion of NADH shuttle systems (Fig. 4) and GPD1 overexpression (Fig. 5A) and which correlates with ChREBP activation (Fig. 5B), binds the GSM with a *K*_d_ value of 17.0 +/- 1.1 μM. Thus, biophysical, genetic, metabolomic and cellular reconstitution data indicate that the GSM of ChREBP should be termed a G3P-sensing module that drives transcription of ChREBPβ, FGF21 and other ChREBP target genes. Notably, this model can account for PPARα-independent induction of FGF21 in conditions of lipolysis^29^, for fructose and glycerol as activators of ChREBP^59^, and for ethanol^45^, reductive stress^46^, hyperglycemia^42^, and mitochondrial dysfunction^43^ as drivers of FGF21 transcription.

**Table 2.** The GSM of ChREBPα directly binds G3P.

| ligand | $K_d$ +/- SD (μM) |
| --- | --- |
| G3P | 17.0 +/- 1.1 |
| G6P | 64.3 +/- 13.5 |
| F6P | 131. +/- 43. |
| X5P | 164. +/- 16. |
| GA3P | 221. +/- 62. |
| F1,6BP | 250. +/- 37. |
| Glc | 270. +/- 54. |
| DHAP | 453. +/- 31. |

After this work was available in preprint form^67^, support for G3P as the activator of ChREBP was provided in an experiment in which reporter plasmid transcription from ChREBP and MLX was reconstituted in HEK293T cells and shown to increase in response to GPD1 overexpression^68^ as we showed in Fig. 5A. Moreover, these investigators added G3P to HEK293T cells expressing ChREBP and MLX to show that G3P addition protected ChREBP from thermal precipitation^68^. As all models of ChREBP activation depend on new protein interactions with small molecule engagement^48,49^, the cellular thermal denaturation assay does constitute evidence for direct binding of G3P. However, the work supports our physiological, metabolomic and biophysical evidence for G3P as the activator of ChREBP.

### The G3P-ChREBP activation system suggests new mechanistic components of the Randle Cycle and cooperation between ChREBP and other transcription factors

Philip Randle’s medical school lectures, published in 1963, contained two simple sketches that depicted what he termed the glucose-fatty acid cycle^69^. The first sketch schematized Glc as the source of G3P, which is the backbone for triglyceride synthesis through the Kennedy pathway^70^. The second sketch illustrated Randle’s observation that in conditions of high fatty acid availability, there are mechanisms to block glycolysis and that, in conditions of high Glc availability, there are mechanisms to block fatty acid oxidation. In subsequent years, the mechanisms for fatty acid oxidation blocking glycolysis were shown to be mediated by citrate as an inhibitor of Glc transport, hexokinase activity and the activity of phosphofructokinase 1^71^.

Regulation in what is termed the sweet side of the Randle Cycle has been explained by the function of malonyl-coA as an essential substrate of fatty acid synthase and inhibitor of carnitine palmitoyltransferase 1, which is required for long chain fatty acid (LCFA) entry into mitochondria^72^. Thus, when acetyl coA carboxylase converts cytosolic Ac-coA to malonyl-coA, it is not only committing carbon flow to synthesis of LCFA but also blocking LCFA oxidation.

While Randle’s principles of fuel utilization have been important in guiding research and medicine, there are unsolved problems in metabolism that remain elusive. For example, though fructose is known to be more lipogenic than Glc, it is not clear that this is fully explained on the basis of the higher affinity fructokinase or bypass of phosphofructokinase regulation^73^.

According to our model, ChREBP evolved specifically to respond to formation of G3P by stimulating transcription of enzymes that convert carbohydrates to LCFAs and stimulating transcription of Kennedy pathway enzymes that linked LCFAs to G3P in triglyceride synthesis. Fructose would thus be more lipogenic than Glc because it is a more potent ChREBP activator^59^ and it is a more potent ChREBP activator because fructolysis (Fig. 1C) would tend to produce more G3P than glycolysis. Notably, because G3P is not a direct glycolytic intermediate like DHAP or GA3P, but rather an electron carrier in the GPDS, production of G3P from carbohydrates is a signal of carbohydrate overload from diet, diabetes, *GCKR* variants, or a signal that would be generated at normoglycemia by fructose, ethanol or mitochondrial insufficiency. Reductive stress was previously noted as a shared mechanism underlying metabolic features of both alcoholic and nonalcoholic hepatic steatosis via ChREBP activation^60^. Identification of G3P as the activator of ChREBP further unites mechanisms of hepatic steatogenesis downstream of ethanol, fructose, hyperglycemia and mitochondrial insufficiency.

We do not suggest that G3P-driven ChREBP activated transcription is fully responsible for complex metabolic switches. Lipogenesis requires activation of both ChREBP and sterol response element binding protein 1c (SREBP-1c)^62^, which occurs with depression of the carnitine palmitoyltransferase and beta oxidation systems. In contrast, fasting-induced lipolysis, which is expected to produce G3P and activate ChREBP, also activates PPARα and the beta oxidation program^27–29^. It is thus to be expected that complex interactions between fatty acid ligands, PPAR isoforms, SREBP and other transcription factors modulate metabolic outputs of ChREBP in changing conditions. Two expected differences between G3P formation from Glc, ethanol and mitochondrial insufficiency (Fig. 1B) and G3P formation from lipolysis is that lipolytic G3P formation is expected to require GK activity and to produce PPARα-activating fatty acids (Fig. 1D). Thus, it is interesting to note that GK expression has been shown to drive lipolytic gene expression in mouse liver, though this was attributed to transcriptional activation of SREBP-1c rather than enzymatic activity^74^.

### The mouse CD model shows an integrated stress response (ISR) that may exacerbate urea cycle dysfunction

The integrated stress response (ISR) is an adaptive response to amino acid deprivation that is mediated by phosphorylation of eukaryotic translational initiation factor eIF2α, production of specific transcription factors ATF4 and/or ATF5, and consequent gene expression to restore protein homeostasis^75^. Under conditions of protein limitation, new protein synthesis is focused on resolving amino acid deficiency. For example, FGF21 is an ISR-activated hormone that drives protein ingestion^76^ while SLC3A2 is an ISR-activated transporter that increases cellular amino acid import^77^. Classical ISR-activated enzymes asparagine (Asn) synthetase ASNS^78^ and cystathionine γ-lyase CTH^79^ are induced to produce Asn and cysteine (Cys), respectively.

Mitochondrial defects are known to induce the ISR^80^ via mechanisms that are not completely understood. However, it was shown that complex I inhibition of C2C12 mouse myoblasts induces the ISR by virtue of depressing Asp and Asn synthesis^81^. When amino acids are in limited supply, uncharged tRNAs activate GCN2 protein kinase to phosphorylate eukaryotic translational initiation factor eIF2α^82^. In myoblasts, complex I inhibition-mediated induction of the ISR can be relieved by lowering the NADH/NAD^+^ ratio and by provision of Asp or Ans^81^.

RNAseq data from the mouse model of CD supports Asp limitation as a mechanism by which the ISR can be activated by mitochondrial dysfunction in liver. Further, the data suggest that the ISR may aggravate urea cycle in CD. As shown in Fig. 6A, we noted that *Asns*^78^ is strongly upregulated with deletion of *Slc25a13*. Moreover, with inactivation of both NADH shuttle systems (Fig. 6B), additional components of an ISR were observed including elevation of the *Atf5* transcription factor^83^. Exposure of *Slc25a13 -/- Gpd2* -/- mice to glycerol (Fig. 6C) increased expression of *Slc3a2*^77^ and *Cth*^79^, which are recognized as part of the ISR^84^.

**Fig 6.**
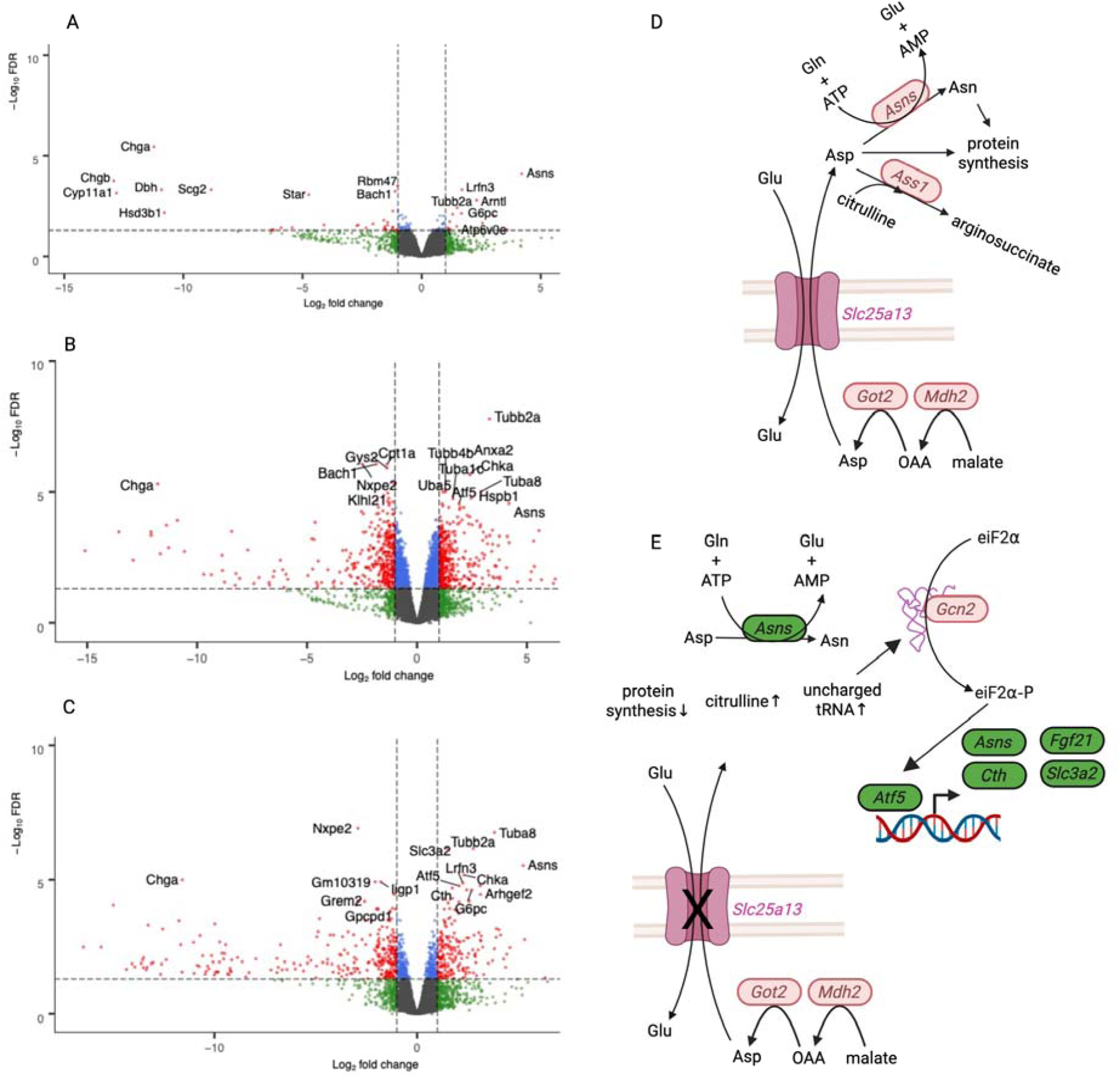
The murine CD model shows evidence of an integrated stress response (ISR) that may underly and aggravate urea cycle dysfunction. Differentially expressed hepatic genes shown in volcano plots for (A) *slc25a13 -/-* mice versus wild-type, (B) *slc25a13 -/- gpd2 -/-* mice versus wild-type and (C) *slc25a13 -/- gpd2 -/-* glycerol-exposed mice versus wild-type water-exposed. In (D), mitochondrial generation of Asp and the role of Slc25a13 in providing Asp to hepatocyte cytosol are depicted. Asp is used by Asn synthetase (Asns) to make Asn, is used in protein synthesis, and is ligated to citrulline by argininosuccinate synthetase 1 (Ass1) in the urea cycle. In (E), an ISR is proposed to be initiated by shortage of cytosolic Asp and the consequent accumulation of uncharged tRNA Asp and/or tRNA Asn, which activate Gcn2 kinase activity on eIF2α. Subsequent induction of Atf5 and Atf5 target genes could aggravate urea cycle dysfunction because Fgf21 drives protein ingestion, Slc3a2 increases amino acid uptake, Asns consumes Asp, further limiting availability of Asp for nitrogen disposal, and Cth produces ammonia likely to exacerbate the citrulline-accumulation problem.

As depicted in Fig. 6D, Asp is produced in mitochondria in a process requiring activities of MDH2, GOT2 and the METC^85^. Asp enters the cytosol through SLC25A13, where it is used for protein synthesis, ASNS-dependent conversion to Asn, and nitrogen disposal through the urea cycle. As in the case of myoblasts treated with piericidin to inhibit complex I, which produced an ISR that was reversed by provision of Asp or Asn^81^, our data show that loss of SLC25A13 is sufficient to induce an ISR. However, whereas the ISR is intended to be an adaptive system to restore protein homeostasis^75^, the genes induced in the model of CD suggest that the ISR may aggravate the ammonia disposal problem for hepatocytes in CD because FGF21 may cause people to eat more protein^76^, SLC3A2 will drive cellular amino acid import into hepatocytes already challenged with nitrogen disposal^77^, ASNS expression would tend to convert urea cycle- limiting levels of Asp for Asn synthesis^78^ and the CTH enzyme activity produces ammonia^79^. Thus, we suggest that partial inhibition of ATF5, and/or target enzymes such as ASNS and CTH could be considered as pharmacological approaches to improve urea cycle function in CD.

### *SLC25A13* mutation carriers have distinct metabolic traits and signs of potential FGF21 overexpression

Given the flux of carbohydrate oxidation, lipolysis, ethanol exposure and other metabolic perturbations that occur over the lifetimes of human beings, we considered whether heterozygosity for *SLC25A13* disease mutation might be associated with anthropometrics, dietary patterns or unusual biomarkers. We used gnomAD^86^ to assemble a list of the most commonly occurring *SLC25A13* alterations that are scored as pathological or likely pathological (Supplemental Table 1) and subjected them to several tests of genetic association. In deep phenotype genome-wide association studies (GWAS), rs80338722 was moderately associated with body weight and BMI in BioBank Japan (*P*=8.3e^-6^ and *P*=4.1e^-4^, respectively)^87^ and associated with triglyceride levels in non-diabetic individuals^88^, among other traits. Three additional alterations of *SLC25A13* variants were associated with total cholesterol^88^, leptin^88^, and autoimmune hepatitis^87^. Our interpretation of these data is that at a population level, *SLC25A13* heterozygosity may bias liver metabolism to produce greater G3P-ChREBP-dependent transcriptional output, which could either predispose to lipogenesis and/or produce a signal for elevated FGF21 as reported^89^ for the *GCKR* polymorphism^44,46^. Indeed, based on known biology^90^ and Mendelian randomization^91^, one might expect higher FGF21 to be associated with higher sodium clearance from kidney and with dietary preference for fatty fish versus sweets^76^. As shown in Fig. 7A, a phenome-wide association study for rs80338722 within BioBank Japan showed an association with low body weight. When we aggregated rare variant gene-based burden tests between *SLC25A13* alterations and 189 complex traits from the Common Metabolic Disease Knowledge Portal, the data suggest a distinct metabolic profile associated with low C-peptide and low incidence of type 1 diabetes but elevated levels of bilirubin and apolipoprotein B, and a strong signal for high urinary sodium excretion (Fig. 7B). Consistent with the possibility that inactivation of one copy of the *SLC25A13* gene elevates FGF21, common variant gene-based tests for *SLC25A13* and 189 complex traits from the Common Metabolic Disease Knowledge Portal^88^ identified a strong association with oily fish consumption (Fig. 7C).

**Fig. 7.**
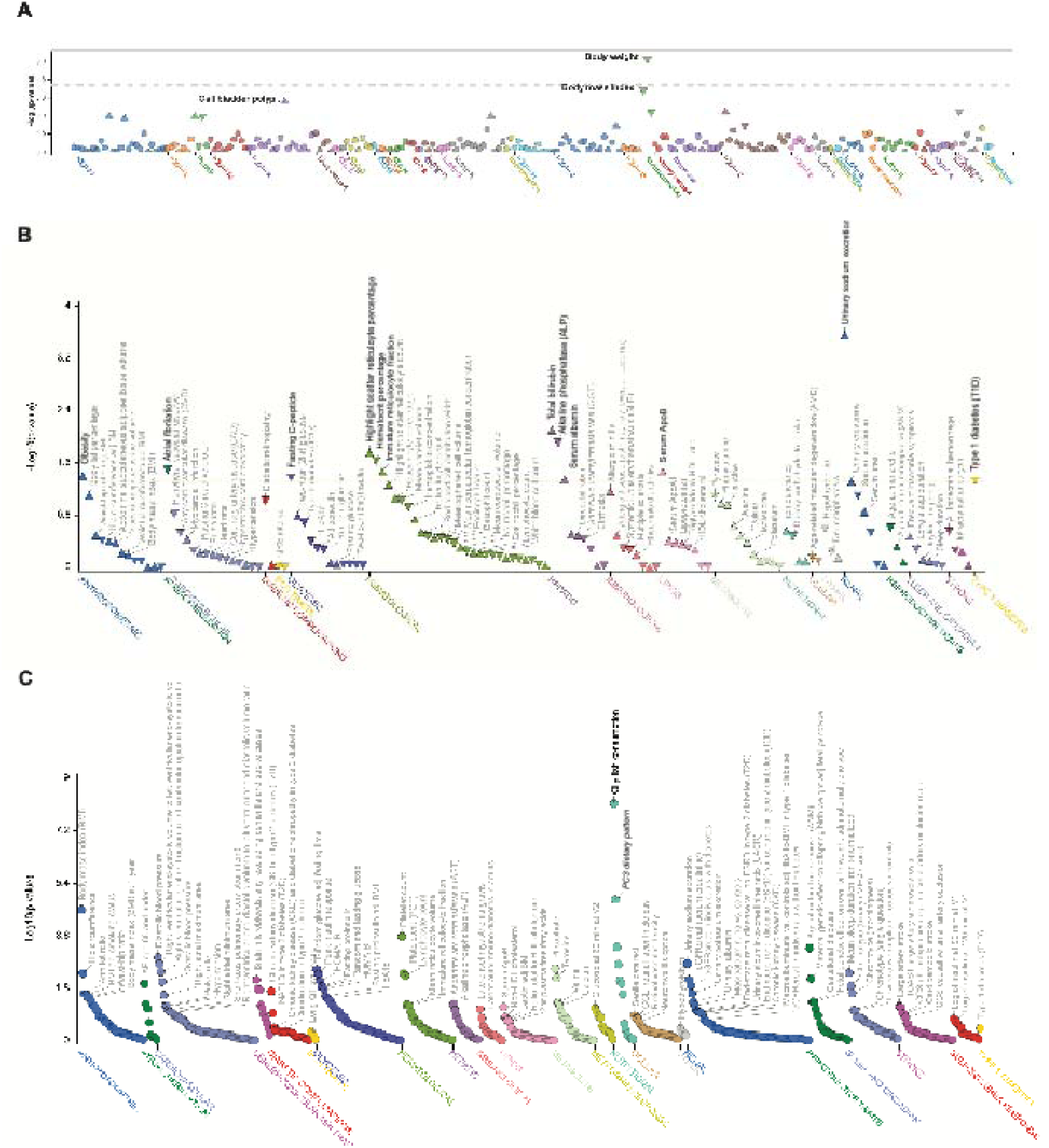
Locus-wide and gene-based association plots for *SLC25A13*. A) A phenome-wide association study for rs80338722 within BioBank Japan identified an association with low body weight. The x-axis represents phenotypes within Biobank Japan, while the y-axis is the -log_10_P- value for association between rs80338722 and each trait. The dashed grey line represents the moderate significance threshold (P < 1e-3). B) Aggregated rare variant gene-based burden tests between *SLC25A13* and 189 complex traits from the Common Metabolic Disease Knowledge Portal identified phenotypic associations, including high urinary sodium excretion and elevated bilirubin. Traits above the orange dashed line (significance threshold P < 5e-2) are statistically significant. Triangles pointing up indicate associations with increased trait levels or disease risk, while triangles pointing down indicated associations with decreased traits levels or disease risk. C) Common variant gene-based tests for *SLC25A13* and 189 complex traits from the Common Metabolic Disease Knowledge Portal identified a strong association with oily fish consumption. Traits above the orange dashed line (threshold P < 2.5e-6) are nominally significant.

Data on human variation suggest that pathological *SLC25A13* mutations appeared multiple times in the Far East^15^. Our data suggest that loss-of-function allele persistence may be mediated by beneficial effects of FGF21 expression on renal function, insulin sensitivity and food choices.

Indeed, if it is true that *SLC25A13* mutation carriers circulate higher levels of FGF21, one might expect that lower ethanol and/or fructose consumption would protect their livers from the potentially lipogenic combination of these energy inputs with diminished flux through the MAS. Specifically, one might predict that *SLC25A13* mutation carriers drink less alcohol than the general population but that those mutation carriers who drink have a greater degree of hepatic steatosis.

### Conclusions

Just as rare mutations in model organisms have revealed complexities of morphogenesis and gene regulation, many fundamental biological insights have been revealed from rare human diseases. In the case of CD, it became apparent that without a key component of the MAS, people and mice are sensitive to development of MASLD despite being lean, and do not enjoy sweets despite an intact detection and initial preference for sweets. We propose that a key to both of these CD presentations is accumulation of hepatic G3P. The G3P-ChREBP program drives a lipogenic transcriptional program that induces *Pklr*, *Acly*, *Acaca*, *Fasn*, *Elovl6* and other genes to synthesize LCFAs, and *Agpat2* and other genes to link newly synthesized LCFAs to the G3P backbone for triglyceride synthesis. Notably, the product of *Acaca*, malonyl-coA, is not only the key substrate for LCFA synthesis but also the key inhibitor of LCFA entry into mitochondria for β-oxidation^72^. Thus, G3P activation of ChREBP has the potential to transcriptionally direct triglyceride synthesis causing hepatic steatogenesis while also promoting resistance to hepatic lipolysis. Indeed, based on the low ATP state of the liver in CD models^21^, one might have expected brisk usage of stored hepatic triglycerides during the fasting daytime of CD mice or the fasting nighttime of patients with CD. However, the fact that lean MASLD is common in CD^18^ and the clinical observation that MCTs are preferable to common fats^5^ suggest that CPT1A may be inhibited in CD, thereby rendering stored triglycerides to be resistant to oxidation.

Additionally, at the RNA level, our data show that the *Slc25a13 -/- Gpd2 -/-* model significantly depresses expression of *Cpt1a* (Fig. 6B). Thus, CD patients and others with MASLD driven by the proposed G3P-ChREBP lipogenesis program may benefit from small molecule activators of CPT1A^92^ or AMP kinase^93^, which would be expected to turn off Ac-coA carboxylase and thereby relieve CPT1A inhibition.

It is notable that the FGF21 induction system responds to a number of conditions of metabolic stress including fasting, and ingestion of fructose and ethanol. While FGF21 signals to the brain to limit sweets and alcohol and to eat protein^76^, it also makes complex signals to the periphery, which have the potential to treat MASLD^94^ and improve renal function^95^. Discovery of the G3P- ChREBP induction system suggests strategies to develop lipidated prodrugs of G3P that would induce FGF21 expression, potentially in combination with fibrates to activate PPARα-dependent lipolysis and synergistically superinduce FGF21^27,29^. While chronic FGF21-elevated conditions such as mitochondrial disease are potentially FGF21-resistant^43^, G3P-releasing FGF21-inducing prodrugs that increase energy expenditure and alter food choices could be valuable to address overweight and MASLD particularly if combined with glucagon-like peptide-1 receptor agonists^96^.

Though ChREBP has long been known to connect carbohydrate oxidation to lipogenesis^48–50^, and G3P has long been known to serve as the backbone for triglyceride synthesis^70^ with a unique location in the glucose-fatty acid cycle^69^, the accumulation of G3P and induction of a G3P- ChREBP transcriptional program in the mouse model of CD allowed us to propose that the most distinctive presentations of CD, namely sweet aversion, lean MASLD and the beneficial effects of MCTs in CD are due to G3P-ChREBP signaling. Ongoing work will further probe components of FGF21 transcriptional induction mechanisms and determine how the ISR and ChREBP induction systems interact in health and disease.

## Acknowledgments

We thank Takeyori Saheki and the Citrin Foundation for donation and shipment of *Slc25a13 -/- Gpd2 -/-* mouse sperm, Walter Tsark at City of Hope for *in vitro* fertilization, and Tsui-Fen Chou at the Proteome Exploration Laboratory of California Institute of Technology for assistance with mass spectrometry. We thank Patrick Fueger, Conn Mallet, John Williams, Zhao Wang, Yingfeng Deng, Sarah Shuck and Rama Natarajan for helpful consultations. Figs. 1 and 6 were made with Biorender.

## Data and materials availability

Mouse data are archived at Dataverse, metabolomic data are at the Metabolomics Workbench, and gene expression data are at the Gene Expression Omnibus. Plasmids have been deposited at Addgene.

## Supplementary Materials

Supplementary Table 1

## Materials and methods

### Generation of mice and mouse samples

All mouse breeding and experiments were performed with protocols approved by the City of Hope Institutional Animal Care and Use Committee and Institutional Biosafety Committee. Sperm from a male *Slc25a13 -/- Gpd2 -/-* C57BL6/J mouse^14^ were a kind gift of Dr. Saheki and the Citrin Foundation. In vitro fertilization was performed with C57BL6/J eggs and a pseudopregnant C57BL6/J female recipient by Walter Tsark in the City of Hope Transgenic Mouse Core. Resulting diheterozygous offspring were intercrossed to generate wild-type, *Slc25a13 -/-*, *Gpd2 -/-*, and *Slc25a13 -/- Gpd2 -/-* mice of both sexes. At weaning, genotypes were determined by PCR of tail tissues (Transnetyx, Cordova, TN), and mice were single-sex group-housed until experimental use. Mice were maintained at 21 °C under a standard 12:12-hour light-dark cycle and provided with *ad libitim* access to food and water. For analysis of FGF21 expression, hepatic gene expression and metabolite accumulation, 8-16 week old mice were single-housed with chow (LabDiet irradiated PicoLab Rodent Diet 20, 5053) and a single bottle of either water or 5% (w/v) glycerol for two days. Mice were then euthanized by decapitation via guillotine without sedation, exsanguinated with a funnel for blood collection into 1.5 ml microcentrifuge tubes on ice, and livers were freeze-clamped in liquid nitrogen immediately after harvest^98,99^. Sera were obtained by centrifuging blood at 8,000 *g* for 10 minutes at 4°C. Liver samples were pulverized with a mortar and pestle cooled by liquid nitrogen.

### FGF21 quantification

Mouse/Rat FGF21 Quantikine ELISA Kit from R&D Systems (Cat. No. MF2100) was used for FGF21 quantification of mouse sera. 50 μl of assay diluent RD1-41 was added to each well followed by equal volumes of standard, control, and experimental samples.

After incubation for 2 hr at room temperature, wells were aspirated and washed 5 times with 400 μl of wash buffer. 100 μl of horseradish peroxidase-conjugated FGF21 antibody solution was then added to each well and incubated at room temperature for 2 hr followed by five washes with 400 μl of wash buffer. After washing, 100 μl of freshly prepared substrate was added to each well and incubated for 30 minutes at room temperature. Assays were terminated with addition of 100 μl of stop solution followed by reading at 450 nm with wavelength correction at 570 nm.

### RNAseq

20 mg liver aliquots were used for RNA isolation (Qiagen, RNeasy Mini Kit, 74104) and DNase I treatment (Qiagen, RNase-Free DNase Set, 79254). RNA integrity was assessed using an Agilent 4200 Tapestation, ensuring RNA integrity number > 5.0 for all samples. Library preparation and sequencing was performed by 7 Traits of Watertown, MA. Libraries were prepared using the NEBNext UltraExpress RNA Library Prep Kit (New England Biolabs, Ipswich, MA), following the manufacturer’s protocol using 150 ng RNA per sample. mRNA was enriched using poly(A) selection. Enriched mRNA was fragmented, reverse transcribed, and converted to cDNA. cDNA underwent end repair, adapter ligation, and size selection. PCR amplification was performed using 11 cycles to ensure sufficient library yield while minimizing amplification bias. Library quality and size distribution were assessed using a Tapestation (Agilent) and quantified by Qubit (Thermo Fisher Scientific). Libraries were pooled on an equimolar basis and sequenced on an Illumina NovaSeq X Plus platform using a 25B lane configuration with paired-end 150 bp reads with a minimum of 20 million paired reads per sample. Base calling and demultiplexing were performed using Illumina’s DRAGEN Bio-IT platform. Raw sequencing reads were processed for quality control using FastQC and trimmed to remove adapters and low-quality bases using Trimmomatic^100^and polyA tails using FASTP^101^.

Processed reads were mapped back to the mouse genome (mm10) using STAR software (v. 2.6.0.a)^102^. HTSeq software (v.0.11.1) was employed to generate the count matrix with default parameters^103^. Differential expression analysis was performed by normalizing read counts to expression values using the TMM normalization method in edgeR^104,105^. Generalized linear models were applied to identify differentially expressed genes between glycerol and water-exposed liver samples of the same genotype or between liver samples of different genotypes. Normalized expression levels from TMM were used as the dependent variable, while sequencing batches were included as an independent variable to account for batch effects. Genes with an FDR-adjusted p-value below 0.05 and a fold change greater than 2 or less than 0.5 were classified as significantly upregulated or downregulated, respectively.

### Metabolomic analysis of mouse livers

For G3P quantification, 2.0 mg samples of pulverized frozen liver were spiked with ^13^C_3_-G3P (2.28 µM final concentration, Sigma Aldrich). Frozen samples were immediately processed with a boiling buffered ethanol extraction (600 µL of 25% 10 mM HEPES Buffer and 75% Ethanol) followed by vigorous vortexing. Samples were placed on a thermomixer for 5 min at 55 °C with shaking at 1,200 rpm, then cooled on ice for 30 seconds. Samples were then placed in a water bath sonicator for 1 minute followed by an additional 30 seconds on ice. Samples were clarified by centrifugation at 16,100 *g* in a prechilled centrifuge at 4 °C; the supernatant was transferred to a clean tube followed by a second round of centrifugation. Clarified supernatants were transferred to new tubes and dried for 5 hours in a vacuum centrifuge at 4 °C. Dried samples were resuspended in 80 µl of LC-MS water. Twenty- folded diluted samples were transferred to MS vials for analysis on LC-MS/MS. For analysis of hexose phosphates and pentose phosphates, 2.0 mg samples of pulverized frozen liver were spiked with a ^13^C glucose-grown yeast extract (ISO101, Cambridge Isotope Laboratories) as an internal standard^106^ and the same workup was performed. Samples were analyzed on a Vanquish Horizon UHPLC with a tandem Thermo Scientific Orbitrap Fusion mass spectrometer. Vials were maintained in autosampler at 4 °C. Instrument source parameters were held at 3 kV negative ion spray voltage, 300°C ion transfer tube temperature, 250 °C vaporizer temperature, sheath gas of 20, auxiliary gas of 10, and sweep gas of 3. Liquid chromatography separation was carried out using an Acquity Premier HSS T3 column with VanGuard FIT, 1.8 µm, 2.1 x 150 mm with mobile phases A (5 mM tributylamine and 10 mM acetic acid with 5% v/v methanol in LC-MS grade water) and B (LC-MS grade methanol) at a constant flow rate of 0.5 ml/min.

Separation was carried out with starting condition of 0% B; 0-10 min, 10.5% B; 10-18 min, 52.6% B; 18-19 min 52.6% B; 19-20 min, 0% B; 20-26 min, 0% B. Spectra for G3P were acquired using a targeted MS2 scan with parent ion of *m/z* 171.0058 with collision energy 25. Spectra for ^13^C_3_-G3P were acquired with parent ion *m/z* of 174.0165 with collision energy 25 with primary fragment ion of *m/z* 78.9588. Hexose phosphate spectra were acquired using a targeted MS2 scan of parent ion *m/z* 259.0198. ^13^C_6_-hexose phosphate spectra were acquired with a parent ion of *m/z* 265.0426 with collision energy of 20 and primary fragment ion *m/z* 96.9690. Pentose phosphate spectra were acquired using targeted MS2 scan for the parent ion *m/z* 229.0124 at collision energy 30 with a primary fragment ion *m/z* of 78.9588.

### Cellular reconstitution of ChREBP-MLX-dependent ChORE-luciferase activity

Human ChREBPα (accession number NM_032951.3), human ChREBPβ (accession number XM_047420437.1), human MLX (accession number NM_170607.3), and eGFP coding sequences were synthesized by Genewiz (Suzhou, China) and cloned into pcDNA3.1 vectors (Invitrogen). GK was subcloned into pcDNA3.1 (Invitrogen) from pWZL-Neo-Myr-Flag-GK (Addgene plasmid 20493). pcDNA3.1-GPD1 and pcDNA3.1-GAPDH were purchased from GenScript (clone ID: OHu20325 and OHu20566). pcDNA3.1-*Lb*NOX, pcDNA3.1-*Ec*STH, pGL4.14 [luc2/Hygro]-ChoRE, and pGL4.75[hRluc/CMV] were as described^60^. HEK293T cells were seeded into 24-well plates and maintained in DMEM medium (Gibco) with 10% FBS (Gibco) and 1% Pen-Strep (Gibco) at 37 °C and 5% CO_2_ overnight. On the following day, media were replaced by OPTI-MEM (Gibco) 1 hour before plasmid co-transfection. Lipofectamine 3000 reagent (Invitrogen) was used for co-transfections for 5 hours, after which media were replaced with DMEM containing 10% FBS for 48 hours. Cells were then collected, and luciferase assays were performed using the Firefly and Renilla Single Tube Luciferase Assay Kit (Biotium) following the manufacturer’s protocol. A Tecan Infinite M Plex plate reader was used to measure Firefly and Renilla luminescence. Firefly luciferase values were normalized to corresponding Renilla luciferase measurements to correct for cell quantity and transfection efficiency.

### Metabolomic analysis of transfected HEK293T cells

HEK293T cells were seeded into 6-well plates overnight and then underwent plasmid co-transfection as described above for 48 hours. Cells were then washed with PBS, quenched with dry ice-cold 80% methanol, and transferred to conical tubes. Relative quantification of 137 metabolites was performed as described^60^.

### Biophysical characterization of the ChREBP GSM

The coding sequence for mouse ChREBPα (amino acids 43 to 307) were converted to *E. coli*-optimized codons and inserted into a pET vector carboxyl to MetHis_6_. The plasmid, pVB240306, was transformed into in *E. coli* Arctic Express, and expression was induced at 12.5 °C with 1 mM IPTG for 18 hrs. Total protein from 1 liter culture was solubilized in 15 ml denaturing lysis buffer (8M urea, 100 mM NaH_2_PO_4_, 10 mM Tris Cl, pH 8.0, 0.05% Tween 20). Recombinant protein was captured by overnight incubation with 3 ml Ni-NTA resin (Qiagen) followed by washing with 30 ml denaturing buffer (8 M urea, 100 mM NaH_2_PO_4_, 10 mM Tris Cl, pH 6.3, 0.05% Tween 20). On-column protein refolding was performed by washing with 30 ml of 50 mM NaH_2_PO_4_, pH 8.0, 300 mM NaCl, 0.1% Triton X-100, followed 30 ml of 50 mM NaH_2_PO_4_, pH 8.0, 300 mM NaCl, 5 mM β-cyclodextrin, 0.05% Tween 20, and 30 ml of 50 mM NaH_2_PO_4_, pH 8.0, 500 mM NaCl, 0.05% Tween 20. His-tagged ChREBP43-307 was recovered in 50 mM NaH_2_PO_4_, pH 8.0, 300 mM NaCl, 0.05% Tween 20, 350 mM imidazole at a yield of 1-2 mg/liter culture. All ligands were prepared in the same buffer. Ligand binding was determined by isothermal titration calorimetry using a Nano ITC (TA Instruments, Waters, USA) to measure ligand concentration-dependent heat changes. Ligands were delivered in a 50 µl syringe to 185 µl His-tagged ChREBP43-307 in the sample cell. After an initial 0.8 µl injection, 19 additional 2.5 µl injections were made with a time interval of 200 s between each injection. Measurements were made at 20 °C with stirring at 250 rpm. Informative ligand concentrations were determined empirically to observe saturable binding. Binding profiles were fitted to a blank and independent model with single-site binding using analysis software supplied with the instrument. Each experiment was performed in triplicate.

## References

1 Palmieri, L. et al. Citrin and aralar1 are Ca(2+)-stimulated aspartate/glutamate transporters in mitochondria. EMBO J 20, 5060–5069, doi:10.1093/emboj/20.18.5060 (2001).

2 Saheki, T. & Kobayashi, K. Mitochondrial aspartate glutamate carrier (citrin) deficiency as the cause of adult-onset type II citrullinemia (CTLN2) and idiopathic neonatal hepatitis (NICCD). J Hum Genet 47, 333–341, doi:10.1007/s100380200046 (2002).

3 Saheki, T. et al. Metabolic derangements in deficiency of citrin, a liver-type mitochondrial aspartate-glutamate carrier. Hepatol Res 33, 181–184, doi:10.1016/j.hepres.2005.09.031 (2005).

4 Grunert, S. C. et al. Citrin deficiency mimicking mitochondrial depletion syndrome. BMC Pediatr 20, 518, doi:10.1186/s12887-020-02409-x (2020).

5 Saheki, T., Moriyama, M., Funahashi, A. & Kuroda, E. AGC2 (Citrin) Deficiency-From Recognition of the Disease till Construction of Therapeutic Procedures. Biomolecules 10, doi:10.3390/biom10081100 (2020).

6 Lin, Y. et al. Combining newborn metabolic and genetic screening for neonatal intrahepatic cholestasis caused by citrin deficiency. J Inherit Metab Dis 43, 467–477, doi:10.1002/jimd.12206 (2020).

7 Kikuchi, A. et al. Simple and rapid genetic testing for citrin deficiency by screening 11 prevalent mutations in SLC25A13. Mol Genet Metab 105, 553–558, doi:10.1016/j.ymgme.2011.12.024 (2012).

8 Kido, J., Makris, G., Santra, S. & Haberle, J. Clinical landscape of citrin deficiency: A global perspective on a multifaceted condition. J Inherit Metab Dis 47, 1144–1156, doi:10.1002/jimd.12722 (2024).

9 Kim, S. Y. & Yi, D. Y. Components of human breast milk: from macronutrient to microbiome and microRNA. Clin Exp Pediatr 63, 301–309, doi:10.3345/cep.2020.00059 (2020).

10 Belenky, P., Bogan, K. L. & Brenner, C. NAD+ metabolism in health and disease. Trends in Biochemical Sciences 32, 12–19, doi:10.1016/j.tibs.2006.11.006 (2007).

11 Bricker, D. K. et al. A mitochondrial pyruvate carrier required for pyruvate uptake in yeast, Drosophila, and humans. Science 337, 96–100, doi:science.1218099 [pii] 10.1126/science.1218099 (2012).

12 Goyal, S. et al. Dynamics of SLC25A51 reveal preference for oxidized NAD(+) and substrate led transport. EMBO Rep 24, e56596, doi:10.15252/embr.202256596 (2023).

13 Dawson, A. G. Oxidation of cytosolic NADH formed during aerobic metabolism in mammalian cells. Trends Biochem Sci 4, 171–176 (1979).

14 Saheki, T. et al. Citrin/mitochondrial glycerol-3-phosphate dehydrogenase double knock- out mice recapitulate features of human citrin deficiency. J Biol Chem 282, 25041–25052, doi:10.1074/jbc.M702031200 (2007).

15 Tavoulari, S., Lacabanne, D., Thangaratnarajah, C. & Kunji, E. R. S. Pathogenic variants of the mitochondrial aspartate/glutamate carrier causing citrin deficiency. Trends Endocrinol Metab 33, 539–553, doi:10.1016/j.tem.2022.05.002 (2022).

16 Gonzalez-Moreno, L. et al. Exogenous aralar/slc25a12 can replace citrin/slc25a13 as malate aspartate shuttle component in liver. Mol Genet Metab Rep 35, 100967, doi:10.1016/j.ymgmr.2023.100967 (2023).

17 Saheki, T. et al. Reduced carbohydrate intake in citrin-deficient subjects. J Inherit Metab Dis 31, 386–394, doi:10.1007/s10545-008-0752-x (2008).

18 Komatsu, M. et al. Citrin deficiency as a cause of chronic liver disorder mimicking non-alcoholic fatty liver disease. J Hepatol 49, 810–820, doi:10.1016/j.jhep.2008.05.016 (2008).

19 Fukushima, K. et al. Conventional diet therapy for hyperammonemia is risky in the treatment of hepatic encephalopathy associated with citrin deficiency. Intern Med 49, 243–247, doi:10.2169/internalmedicine.49.2712 (2010).

20 Sinasac, D. S. et al. Slc25a13-knockout mice harbor metabolic deficits but fail to display hallmarks of adult-onset type II citrullinemia. Mol Cell Biol 24, 527–536, doi:10.1128/MCB.24.2.527-536.2004 (2004).

21 Saheki, T. et al. Oral aversion to dietary sugar, ethanol and glycerol correlates with alterations in specific hepatic metabolites in a mouse model of human citrin deficiency. Mol Genet Metab 120, 306–316, doi:10.1016/j.ymgme.2017.02.004 (2017).

22 Flippo, K. H. & Potthoff, M. J. Metabolic Messengers: FGF21. Nat Metab 3, 309–317, doi:10.1038/s42255-021-00354-2 (2021).

23 Jensen-Cody, S. O. et al. FGF21 Signals to Glutamatergic Neurons in the Ventromedial Hypothalamus to Suppress Carbohydrate Intake. Cell Metab 32, 273–286 e276, doi:10.1016/j.cmet.2020.06.008 (2020).

24 Flippo, K. H. et al. FGF21 suppresses alcohol consumption through an amygdalo-striatal circuit. Cell Metab 34, 317–328 e316, doi:10.1016/j.cmet.2021.12.024 (2022).

25 Xu, J. et al. Fibroblast growth factor 21 reverses hepatic steatosis, increases energy expenditure, and improves insulin sensitivity in diet-induced obese mice. Diabetes 58, 250–259, doi:10.2337/db08-0392 (2009).

26 Zouhar, P. et al. A pyrexic effect of FGF21 independent of energy expenditure and UCP1. Mol Metab 53, 101324, doi:10.1016/j.molmet.2021.101324 (2021).

27 Inagaki, T. et al. Endocrine regulation of the fasting response by PPARalpha-mediated induction of fibroblast growth factor 21. Cell Metab 5, 415–425, doi:10.1016/j.cmet.2007.05.003 (2007).

28 Potthoff, M. J. et al. FGF21 induces PGC-1alpha and regulates carbohydrate and fatty acid metabolism during the adaptive starvation response. Proc Natl Acad Sci USA 106, 10853–10858, doi:10.1073/pnas.0904187106 (2009).

29 Badman, M. K. et al. Hepatic fibroblast growth factor 21 is regulated by PPARalpha and is a key mediator of hepatic lipid metabolism in ketotic states. Cell Metab 5, 426–437, doi:10.1016/j.cmet.2007.05.002 (2007).

30 Forman, B. M., Chen, J. & Evans, R. M. Hypolipidemic drugs, polyunsaturated fatty acids, and eicosanoids are ligands for peroxisome proliferator-activated receptors alpha and delta. Proc Natl Acad Sci U S A 94, 4312–4317, doi:10.1073/pnas.94.9.4312 (1997).

31 Uebanso, T. et al. Paradoxical regulation of human FGF21 by both fasting and feeding signals: is FGF21 a nutritional adaptation factor? PLoS One 6, e22976, doi:10.1371/journal.pone.0022976 (2011).

32 Soberg, S. et al. FGF21 Is a Sugar-Induced Hormone Associated with Sweet Intake and Preference in Humans. Cell Metab 25, 1045–1053 e1046, doi:10.1016/j.cmet.2017.04.009 (2017).

33 von Holstein-Rathlou, S. et al. FGF21 Mediates Endocrine Control of Simple Sugar Intake and Sweet Taste Preference by the Liver. Cell Metab 23, 335–343, doi:10.1016/j.cmet.2015.12.003 (2016).

34 Iroz, A. et al. A Specific ChREBP and PPARalpha Cross-Talk Is Required for the Glucose-Mediated FGF21 Response. Cell Rep 21, 403–416, doi:10.1016/j.celrep.2017.09.065 (2017).

35 Dushay, J. R. et al. Fructose ingestion acutely stimulates circulating FGF21 levels in humans. Mol Metab 4, 51–57, doi:10.1016/j.molmet.2014.09.008 (2015).

36 Fisher, F. M. et al. A critical role for ChREBP-mediated FGF21 secretion in hepatic fructose metabolism. Mol Metab 6, 14–21, doi:10.1016/j.molmet.2016.11.008 (2017).

37 Galman, C. et al. The circulating metabolic regulator FGF21 is induced by prolonged fasting and PPARalpha activation in man. Cell Metab 8, 169–174, doi:10.1016/j.cmet.2008.06.014 (2008).

38 Kim, K. H. et al. Acute exercise induces FGF21 expression in mice and in healthy humans. PLoS One 8, e63517, doi:10.1371/journal.pone.0063517 (2013).

39 Cuevas-Ramos, D. et al. Exercise increases serum fibroblast growth factor 21 (FGF21) levels. PLoS One 7, e38022, doi:10.1371/journal.pone.0038022 (2012).

40 Markan, K. R. et al. Circulating FGF21 is liver derived and enhances glucose uptake during refeeding and overfeeding. Diabetes 63, 4057–4063, doi:10.2337/db14-0595 (2014).

41 Zhang, X. et al. Serum FGF21 levels are increased in obesity and are independently associated with the metabolic syndrome in humans. Diabetes 57, 1246–1253, doi:10.2337/db07-1476 (2008).

42 Chen, W. W. et al. Circulating FGF-21 levels in normal subjects and in newly diagnose patients with Type 2 diabetes mellitus. Exp Clin Endocrinol Diabetes 116, 65–68, doi:10.1055/s-2007-985148 (2008).

43 Tyynismaa, H. et al. Mitochondrial myopathy induces a starvation-like response. Hum Mol Genet 19, 3948–3958, doi:10.1093/hmg/ddq310 (2010).

44 Cheung, C. Y. Y. et al. An Exome-Chip Association Analysis in Chinese Subjects Reveals a Functional Missense Variant of GCKR That Regulates FGF21 Levels. Diabetes 66, 1723–1728, doi:10.2337/db16-1384 (2017).

45 Desai, B. N. et al. Fibroblast growth factor 21 (FGF21) is robustly induced by ethanol and has a protective role in ethanol associated liver injury. Mol Metab 6, 1395–1406, doi:10.1016/j.molmet.2017.08.004 (2017).

46 Goodman, R. P. et al. Hepatic NADH reductive stress underlies common variation in metabolic traits. Nature 583, 122–126, doi:10.1038/s41586-020-2337-2 (2020).

47 Laeger, T. et al. FGF21 is an endocrine signal of protein restriction. J Clin Invest 124, 3913–3922, doi:10.1172/JCI74915 (2014).

48 Katz, L. S., Baumel-Alterzon, S., Scott, D. K. & Herman, M. A. Adaptive and maladaptive roles for ChREBP in the liver and pancreatic islets. J Biol Chem 296, 100623, doi:10.1016/j.jbc.2021.100623 (2021).

49 Abdul-Wahed, A., Guilmeau, S. & Postic, C. Sweet Sixteenth for ChREBP: Established Roles and Future Goals. Cell Metab 26, 324–341, doi:10.1016/j.cmet.2017.07.004 (2017).

50 Kawaguchi, T., Osatomi, K., Yamashita, H., Kabashima, T. & Uyeda, K. Mechanism for fatty acid “sparing” effect on glucose-induced transcription: regulation of carbohydrate-responsive element-binding protein by AMP-activated protein kinase. J Biol Chem 277, 3829–3835, doi:10.1074/jbc.M107895200 (2002).

51 Ma, L., Robinson, L. N. & Towle, H. C. ChREBP*Mlx is the principal mediator of glucose-induced gene expression in the liver. J Biol Chem 281, 28721–28730, doi:10.1074/jbc.M601576200 (2006).

52 Herman, M. A. et al. A novel ChREBP isoform in adipose tissue regulates systemic glucose metabolism. Nature 484, 333–338, doi:10.1038/nature10986 (2012).

53 Shih, H. M., Liu, Z. & Towle, H. C. Two CACGTG motifs with proper spacing dictate the carbohydrate regulation of hepatic gene transcription. J Biol Chem 270, 21991–21997, doi:10.1074/jbc.270.37.21991 (1995).

54 Li, M. V., Chang, B., Imamura, M., Poungvarin, N. & Chan, L. Glucose-dependent transcriptional regulation by an evolutionarily conserved glucose-sensing module. Diabetes 55, 1179–1189, doi:10.2337/db05-0822 (2006).

55 McFerrin, L. G. & Atchley, W. R. A novel N-terminal domain may dictate the glucose response of Mondo proteins. PLoS One 7, e34803, doi:10.1371/journal.pone.0034803 (2012).

56 Li, M. V. et al. Glucose-6-phosphate mediates activation of the carbohydrate responsive binding protein (ChREBP). Biochem Biophys Res Commun 395, 395–400, doi:10.1016/j.bbrc.2010.04.028 (2010).

57 Arden, C. et al. Fructose 2,6-bisphosphate is essential for glucose-regulated gene transcription of glucose-6-phosphatase and other ChREBP target genes in hepatocytes. Biochem J 443, 111–123, doi:10.1042/BJ20111280 (2012).

58 Kabashima, T., Kawaguchi, T., Wadzinski, B. E. & Uyeda, K. Xylulose 5-phosphate mediates glucose-induced lipogenesis by xylulose 5-phosphate-activated protein phosphatase in rat liver. Proc Natl Acad Sci U S A 100, 5107–5112, doi:10.1073/pnas.0730817100 (2003).

59 Kim, M. S. et al. ChREBP regulates fructose-induced glucose production independently of insulin signaling. J Clin Invest 126, 4372–4386, doi:10.1172/JCI81993 (2016).

60 Singh, C. et al. ChREBP is activated by reductive stress and mediates GCKR-associated metabolic traits. Cell Metab 36, 144–158, doi:10.1016/j.cmet.2023.11.010 (2024).

61 Kukurba, K. R. & Montgomery, S. B. RNA Sequencing and Analysis. Cold Spring Harb Protoc 2015, 951–969, doi:10.1101/pdb.top084970 (2015).

62 Linden, A. G. et al. Interplay between ChREBP and SREBP-1c coordinates postprandial glycolysis and lipogenesis in livers of mice. J Lipid Res 59, 475–487, doi:10.1194/jlr.M081836 (2018).

63 Li, B. & Dewey, C. N. RSEM: accurate transcript quantification from RNA-Seq data with or without a reference genome. BMC Bioinformatics 12, 323, doi:10.1186/1471-2105-12-323 (2011).

64 Ge, Q. et al. Structural characterization of a unique interface between carbohydrate response element-binding protein (ChREBP) and 14-3-3beta protein. J Biol Chem 287, 41914–41921, doi:10.1074/jbc.M112.418855 (2012).

65 Stoltzman, C. A. et al. Glucose sensing by MondoA:Mlx complexes: a role for hexokinases and direct regulation of thioredoxin-interacting protein expression. Proc Natl Acad Sci U S A 105, 6912–6917, doi:10.1073/pnas.0712199105 (2008).

66 Velazquez-Campoy, A., Ohtaka, H., Nezami, A., Muzammil, S. & Freire, E. Isothermal titration calorimetry. Curr Protoc Cell Biol Chapter 17, Unit 17 18, doi:10.1002/0471143030.cb1708s23 (2004).

67 Tiwari, V. et al. Glycerol-3-phosphate activates ChREBP, FGF21 transcription and lipogenesis in Citrin Deficiency. bioRxiv, 10.1101/2024.12.27.630525 (2024).

68 Cheong, M. C. et al. Ethanol induction of FGF21 in the liver is dependent on histone acetylation and ligand activation of ChREBP by glycerol-3-phosphate. Proc Natl Acad Sci U S A 122, e2505263122, doi:10.1073/pnas.2505263122 (2025).

69 Randle, P. J., Garland, P. B., Hales, C. N. & Newsholme, E. A. The glucose fatty-acid cycle. Its role in insulin sensitivity and the metabolic disturbances of diabetes mellitus. Lancet 1, 785–789, doi:10.1016/s0140-6736(63)91500-9 (1963).

70 Kennedy, E. P. Biosynthesis of complex lipids. Fed Proc 20, 934–940 (1961).

71 Hue, L. & Taegtmeyer, H. The Randle cycle revisited: a new head for an old hat. Am J Physiol Endocrinol Metab 297, E578–591, doi:10.1152/ajpendo.00093.2009 (2009).

72 McGarry, J. D., Leatherman, G. F. & Foster, D. W. Carnitine palmitoyltransferase I. The site of inhibition of hepatic fatty acid oxidation by malonyl-CoA. J Biol Chem 253, 4128–4136 (1978).

73 Softic, S., Cohen, D. E. & Kahn, C. R. Role of Dietary Fructose and Hepatic De Novo Lipogenesis in Fatty Liver Disease. Dig Dis Sci 61, 1282–1293, doi:10.1007/s10620-016-4054-0 (2016).

74 Ouyang, S. et al. Glycerol Kinase Drives Hepatic de novo Lipogenesis and Triglyceride Synthesis in Nonalcoholic Fatty Liver by Activating SREBP-1c Transcription, Upregulating DGAT1/2 Expression, and Promoting Glycerol Metabolism. Adv Sci (Weinh), e2401311, doi:10.1002/advs.202401311 (2024).

75 Costa-Mattioli, M. & Walter, P. The integrated stress response: From mechanism to disease. Science 368, doi:10.1126/science.aat5314 (2020).

76 Khan, M. S. H. et al. FGF21 acts in the brain to drive macronutrient-specific changes in behavioral motivation and brain reward signaling. Mol Metab 91, 102068, doi:10.1016/j.molmet.2024.102068 (2024).

77 de la Ballina, L. R. et al. Amino Acid Transport Associated to Cluster of Differentiation 98 Heavy Chain (CD98hc) Is at the Cross-road of Oxidative Stress and Amino Acid Availability. J Biol Chem 291, 9700–9711, doi:10.1074/jbc.M115.704254 (2016).

78 Lomelino, C. L., Andring, J. T., McKenna, R. & Kilberg, M. S. Asparagine synthetase: Function, structure, and role in disease. J Biol Chem 292, 19952–19958, doi:10.1074/jbc.R117.819060 (2017).

79 Kraus, J. P. et al. Cystathionine gamma-lyase: Clinical, metabolic, genetic, and structural studies. Mol Genet Metab 97, 250–259, doi:10.1016/j.ymgme.2009.04.001 (2009).

80 Nikkanen, J. et al. Mitochondrial DNA Replication Defects Disturb Cellular dNTP Pools and Remodel One-Carbon Metabolism. Cell Metab 23, 635–648, doi:10.1016/j.cmet.2016.01.019 (2016).

81 Mick, E. et al. Distinct mitochondrial defects trigger the integrated stress response depending on the metabolic state of the cell. Elife 9, doi:10.7554/eLife.49178 (2020).

82 Dong, J., Qiu, H., Garcia-Barrio, M., Anderson, J. & Hinnebusch, A. G. Uncharged tRNA activates GCN2 by displacing the protein kinase moiety from a bipartite tRNA-binding domain. Mol Cell 6, 269–279, doi:10.1016/s1097-2765(00)00028-9 (2000).

83 Watatani, Y. et al. Stress-induced translation of ATF5 mRNA is regulated by the 5’- untranslated region. J Biol Chem 283, 2543–2553, doi:10.1074/jbc.M707781200 (2008).

84 Lee, J. I. et al. HepG2/C3A cells respond to cysteine deprivation by induction of the amino acid deprivation/integrated stress response pathway. Physiol Genomics 33, 218–229, doi:10.1152/physiolgenomics.00263.2007 (2008).

85 Birsoy, K. et al. An Essential Role of the Mitochondrial Electron Transport Chain in Cell Proliferation Is to Enable Aspartate Synthesis. Cell 162, 540–551, doi:10.1016/j.cell.2015.07.016 (2015).

86 Gudmundsson, S. et al. Variant interpretation using population databases: Lessons from gnomAD. Hum Mutat 43, 1012–1030, doi:10.1002/humu.24309 (2022).

87 Sakaue, S. et al. A cross-population atlas of genetic associations for 220 human phenotypes. Nat Genet 53, 1415–1424, doi:10.1038/s41588-021-00931-x (2021).

88 Costanzo, M. C. et al. The Type 2 Diabetes Knowledge Portal: An open access genetic resource dedicated to type 2 diabetes and related traits. Cell Metab 35, 695–710 e696, doi:10.1016/j.cmet.2023.03.001 (2023).

89 Larsson, S. C., Michaelsson, K., Mola-Caminal, M., Hoijer, J. & Mantzoros, C. S. Genome-wide association and Mendelian randomization study of fibroblast growth factor 21 reveals causal associations with hyperlipidemia and possibly NASH. Metabolism 137, 155329, doi:10.1016/j.metabol.2022.155329 (2022).

90 Turner, T. et al. FGF21 increases water intake, urine output and blood pressure in rats. PLoS One 13, e0202182, doi:10.1371/journal.pone.0202182 (2018).

91 Giontella, A. et al. Renoprotective effects of genetically proxied fibroblast growth factor 21: Mendelian randomization, proteome-wide and metabolome-wide association study. Metabolism 145, 155616, doi:10.1016/j.metabol.2023.155616 (2023).

92 Liang, K. Mitochondrial CPT1A: Insights into structure, function, and basis for drug development. Front Pharmacol 14, 1160440, doi:10.3389/fphar.2023.1160440 (2023).

93 Srivastava, R. A. et al. AMP-activated protein kinase: an emerging drug target to regulate imbalances in lipid and carbohydrate metabolism to treat cardio-metabolic diseases. J Lipid Res 53, 2490–2514, doi:10.1194/jlr.R025882 (2012).

94 Yang, R., Xu, A. & Kharitonenkov, A. Another Kid on the Block: Long-acting FGF21 Analogue to Treat Dyslipidemia and Fatty Liver. J Clin Endocrinol Metab 107, e417–e419, doi:10.1210/clinem/dgab686 (2022).

95 Li, F., Liu, Z., Tang, C., Cai, J. & Dong, Z. FGF21 is induced in cisplatin nephrotoxicity to protect against kidney tubular cell injury. FASEB J 32, 3423–3433, doi:10.1096/fj.201701316R (2018).

96 Gilroy, C. A. et al. Sustained release of a GLP-1 and FGF21 dual agonist from an injectable depot protects mice from obesity and hyperglycemia. Sci Adv 6, eaaz9890, doi:10.1126/sciadv.aaz9890 (2020).

97 Sillero, M. A., Sillero, A. & Sols, A. Enzymes involved in fructose metabolism in liver and the glyceraldehyde metabolic crossroads. Eur J Biochem 10, 345–350, doi:10.1111/j.1432-1033.1969.tb00696.x (1969).

98 Trammell, S. A. et al. Nicotinamide riboside is uniquely and orally bioavailable in mice and humans. Nat Commun 7, 12948, doi:10.1038/ncomms12948 (2016).

99 Trammell, S. A. et al. Nicotinamide Riboside Opposes Type 2 Diabetes and Neuropathy in Mice. Sci Rep 6, 26933, doi:10.1038/srep26933 (2016).

100 Bolger, A. M., Lohse, M. & Usadel, B. Trimmomatic: a flexible trimmer for Illumina sequence data. Bioinformatics 30, 2114–2120, doi:10.1093/bioinformatics/btu170 (2014).

101 Chen, S., Zhou, Y., Chen, Y. & Gu, J. fastp: an ultra-fast all-in-one FASTQ preprocessor. Bioinformatics 34, i884–i890, doi:10.1093/bioinformatics/bty560 (2018).

102 Dobin, A. et al. STAR: ultrafast universal RNA-seq aligner. Bioinformatics 29, 15–21, doi:10.1093/bioinformatics/bts635 (2013).

103 Anders, S. & Huber, W. Differential expression analysis for sequence count data. Genome Biol 11, R106, doi:10.1186/gb-2010-11-10-r106 (2010).

104 Robinson, M. D., McCarthy, D. J. & Smyth, G. K. edgeR: a Bioconductor package for differential expression analysis of digital gene expression data. Bioinformatics 26, 139–140, doi:10.1093/bioinformatics/btp616 (2010).

105 McCarthy, D. J., Chen, Y. & Smyth, G. K. Differential expression analysis of multifactor RNA-Seq experiments with respect to biological variation. Nucleic Acids Res 40, 4288–4297, doi:10.1093/nar/gks042 (2012).

106 Trammell, S. A. & Brenner, C. Targeted, LCMS-based Metabolomics for Quantitative Measurement of NAD(+) Metabolites. Comput Struct Biotechnol J 4, e201301012, doi:10.5936/csbj.201301012 (2013).

