## Supplementary Table 1 for "Glycerol-3-phosphate activates ChREBP, FGF21 transcription and lipogenesis in Citrin Deficiency"

**Supplementary Table 1. Most frequent pathological variants of *SLC25A13* in gnomAD (March 12, 2024)**

| RSID | Allele count | Origin |
| --- | --- | --- |
| rs80338720 | 185 | Asian |
| rs80338722 | 102 | Asian |
| rs780525233 | 84 | European |
| rs80338716 | 73 | European |
| rs80338729 | 58 | South Asian |
| rs200237622 | 56 | Ashkenazi |
| rs80338721 | 32 | European |
| rs80338725 | 30 | Asian |
| rs781452100 | 30 | European |
| rs80338723 | 23 | Asian |
| rs143181462 | 22 | European |
